# The spatial landscape of glial pathology and T-cell response in Parkinson’s disease substantia nigra

**DOI:** 10.1101/2024.01.08.574736

**Authors:** Kelly Jakubiak, Fahad Paryani, Adithya Kannan, Jaeseung Lee, Nacoya Madden, Juncheng Li, David Chen, Aayushi Mahajan, Shengnan Xia, Xena Flowers, Vilas Menon, David Sulzer, James Goldman, Peter A. Sims, Osama Al-Dalahmah

## Abstract

Parkinson’s Disease (PD) is a progressive neurodegenerative disease that leads to debilitating movement disorders and often dementia. Recent evidence, including identification of specific peripheral T-cell receptor sequences, indicates that the adaptive immune response is associated with disease pathogenesis. However, the properties of T-cells in the brain regions where neurons degenerate are not well characterized. We have analyzed the identities and interactions of T-cells in PD in post-mortem brain tissue using single nucleus RNA sequencing, spatial transcriptomics and T-cell receptor sequencing. We found that T-cells in the substantia nigra of PD brain donors exhibit a CD8+ resident memory phenotype, increased clonal expansion, and altered spatial relationships with astrocytes, myeloid cells, and endothelial cells. We also describe regional differences in astrocytic responses to neurodegeneration. Our findings nominate potential molecular and cellular candidates that allow a deeper understanding of the pathophysiology of neurodegeneration in PD. Together, our work represents a major single nucleus and spatial transcriptional resource for the fields of neurodegeneration and PD.

## Introduction

Parkinson’s disease (PD) is a common neurodegenerative disease, with an incidence exceeded only by Alzheimer’s disease^1^. PD neuropathology is characterized by the aggregates of alpha-synuclein in neurons known as Lewy bodies and Lewy neurites^2^, and a loss of dopaminergic neurons in the substantia nigra (SN)^3^. While current treatments alleviate PD symptoms^4^, they do not slow PD progression, and a better understanding of the disease pathophysiology is needed to identify therapeutic strategies.

Neuropathological studies have previously identified autoimmune features associated with PD, including an increase in T-cell populations in the SN of PD patients^5,6^. T-cells in the peripheral blood recognize and proliferate in response to an alpha-synuclein antigen challenge in PD patients^7^, and an association between neurodegeneration and microglial activation is well-established in other neurodegenerative diseases such as Alzheimer’s^8,9^, but little is known about these phenomena in the PD brain. The brain microenvironment in the PD SN is considered “pro-inflammatory”^10^, and pro-inflammatory microglia may contribute to the pathogenesis and neuronal death in PD^11^. It has also been suggested that microglia are activated in PD by exosomes secreted from neurons with alpha-synuclein aggregates^12^, and astrocytes have been shown to adopt abnormal phenotypes in PD neuropathology that could be associated with antigen presentation pathways^13,14^. Thus, the interaction between brain microenvironment cells and cells of the immune system is worth further investigation.

In animal models, mice that overexpress alpha-synuclein exhibit dopaminergic neurodegeneration following a bout of enteric infection, and this is associated with a substantial entry of peripheral T-cells into the brain^15,16^. A presentation of mitochondrial antigens has also been implicated in adaptive immunity in animal models of PD^17,18^. Finally, T-cells have been shown to adopt reactive phenotypes in PD, and to contribute to neurodegeneration alongside microglia^19,20^.

Together, the clinical and basic data point towards an important role for infiltrating T-cells in the brain during PD pathogenesis. However, previous studies have mainly focused on the characterization of peripheral T-cells in the blood and cerebrospinal fluid^21–27^, leaving the central role of T-cells in the human SN in PD unknown. Additionally, many studies characterizing T-cells of the PD brain rely on immunohistochemistry (IHC) and/or murine data^5,28–31^, and questions of transcriptional profiles of T-cells in the human PD brain remain unanswered. As such, there has also been no effort to compare peripheral and CNS T-cells in PD.

The goal of this study is to create a resource for T-cell and glial pathology in the human postmortem brain. This allows us to characterize the phenotype of the adaptive immune response in human PD brain, and the relationship between central and peripheral T-cells and other cells in the brain microenvironment, mainly focusing on astrocytes and microglia/myeloid cells. To do so, we have analyzed human brain tissue samples from the SN and the cingulate cortex, comparing control and PD. We have used multiple cutting-edge technologies paired with advanced computational techniques, including molecular analysis of one of the highest numbers of PD brain T-cells that have been reported in previous study cohorts^32–36^.

Together, by studying one of the largest human PD T-cell cohorts in the PD brain to date, we present several conclusions. First, we found that T-cells of the PD SN are mainly CD8+, or cytotoxic, and display a tissue resident, clonally expanded phenotype. Second, we find that not only are T-cells increased in the perivascular spaces as previously reported^37^, but also in the brain parenchyma. We also characterized the phenotypes of astrocytes in the SN and cingulate cortex and found marked differences between the two regions. Importantly and unlike the cingulate cortex, PD reactive astrocytes showed decreased *MT3* expression in the SN, a gene we previously showed to be neuroprotective^38^. Also, we employed spatial transcriptomics and described significant changes in the spatial correlation patterns between T-cells and astrocytes. Finally, we performed computational analyses to nominate candidate molecular and cellular interactions that may perpetuate neurodegeneration in PD. Altogether, our results uncover novel insights into the potential roles of glial and T-cell pathology in PD.

## Materials and Methods

### Human Subjects and Brain Tissue

All study protocols were approved by Columbia University Irving Medical Center Institutional Review Board. Postmortem cingulate cortex or SN specimens frozen during autopsy from control (individuals whose brains did not show significant neuropathology) and PD/DLB were obtained from the New York Brain Bank. The tissue was dissected by a board-certified neuropathologist (OAD), or under the supervision of a board-certified neuropathologist. Forty-four cases were selected for snRNAseq and TCR sequencing, each with RNA integrity numbers of >7, and ten of these were selected for spatial transcriptomics analysis. Cortical wedges, excluding subcortical white matter, or SN tissue measuring ∼ 5 x 4 x 3 mm were dissected on a dry ice cooled stage and processed immediately as described below. The demographics of the cases used are provided in **Table S1**.

### TCR Sequencing

To prepare our TCR libraries, we followed the iRepertoire Bulk Reagent Universal User Manual (V20200818). The starting material was 500ng RNA per sample. We used 9 barcodes - HTAIvc kits (HTAIvc01, HTAIvc02, HTAIvc03, HTAIvc04, HTAIvc05, HTAIvc06, HTAIvc07, HTAIvc08 and HTAIvc09). We pooled one library from each of the barcoded kits together for each sequencing run. The libraries were pooled with 10% PhiX spike-in and sequenced with NextSeq High Output 300 Cycles kits (Illumina) on an Illumina NextSeq 550 (read 1: 155 cycles; read 2: 155 cycles). Five total sequencing runs were conducted.

### TCR Data Processing and QC

Raw data processing was performed in accordance with Sims et al., 2016^39^. As such, raw paired-end fastq files were demultiplexed based on the internal 6-nt barcode sequences added during library construction. FLASH 1.2.11 (flash –M 250 –O)^40^ was used to merge the paired reads, which were aligned to the human genome (GRCh37) using the Burrows-Wheeler Aligner (bwa-mem)^41^. Reads mapping to the T-cell receptor loci (TRA) and associated with V- and J-cassettes were extracted and translated in silico in all three readings. Reading frames containing a C…FGXG amino acid motif that was uninterrupted by a stop codon were identified as productive CDR3 amino acid sequences. For each demultiplexed^39^ sample, all V- and J-cassettes were then reference-corrected and the number of reads identified with each unique combination of V- and J-cassettes encoding a CDR3 amino acid sequence were counted. Further, saturation levels for all sample libraries were assessed using the estimate_saturation function from the RNAseQC^42^ package in R, at a depth of 200 and using 10 reps, and saturation curves were plotted for each sample (**Table S2**).

### Calculating Entropy in TCR Data

We used Shannon entropy as a measure of diversity in our TCR dataset^39,43^ for each of the components of the clonotype (CDR3 amino acid sequence, VJ combo, whole clonotype, and the difference between the whole clonotype and the VJ combo), using their reads as a direct measure of their frequency in the repertoire. To do so, the following equation was used:

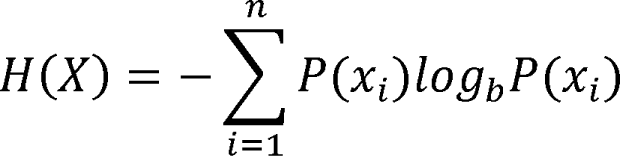

where H is Shannon entropy, X is the clonotype, and the probability of X was calculated from the frequency of reads for the given clonotype X in the entire repertoire of a given sample. We were able to collect entropy values for each sample repertoire, for each clonotype component. We then averaged these entropy values for each region and condition for comparison.

Comparisons between these average values were conducted using linear regression, with formula Entropy ∼ Region + Condition.

### CDR3 Sequence Clustering and Clonal Expansion

To determine which motifs defined the sequences in our TCR repertoire, we used GLIPH2^44^ to extract conserved motifs across CDR3 sequences. We input our CDR3, V, and J sequences and their number of reads into the gliph2 function with default parameters. 16,875 global clusters (one amino acid substitution at a location in a sequence is allowed) and 3,103 local clusters (strings of amino acids with no substitutions) were identified. The clonal expansion metric calculated by GLIPH2 represents the likelihood of a given cluster’s depth being generated by random chance, as measured through permutation testing. To report these values, we subtracted the clonal expansion scores from 1 so that higher clonal expansion scores represent a higher likelihood and lower scores represent a lower likelihood of clonal expansion within a cluster.

To calculate clonal expansion scores for individual subjects, we first filtered our clonotypes by counting the occurrence, or number of reads, of each clonotype. We calculated the mean and variance of each clonotype occurrence across all patients. We then filtered out clonotypes with a mean occurrence less than 1, and variance less than 2. The filtered-out clonotypes served as our background. This was a necessary step for random sampling, as the majority of clonotypes in these repertoires comprised just one or two reads and made it difficult to generate interpretable data.

To generate a quantitative measurement of clonal expansion, we performed random sampling of 1,000 clonotypes from the filtered data and displayed the histogram on a log scale. The histogram revealed two distinct distributions: the background distribution (a right-skewed distribution which contained clonotypes that occur only a handful of times but are abundant by nature), and the signal distribution (a left-skewed distribution which contained clonotypes occurring at far higher frequency and thus more likely to be actively involved in clonal expansion). To extract the signal clonotypes, we required the log frequency of the clonotypes to be greater than 3. We then extracted the signal clonotypes from each patient and generated 100 random distributions as described above and performed a KS test, which generated a D-value that we collected across all patients and grouped based on condition and brain region. We directly compared each subject’s D-value, grouped by brain region and condition, by unpaired t-test. We repeated this 100 times to generate 100 p values to further test the hypothesis of increased clonal expansion. The code used for this analysis can be found at: https://github.com/dalhoomist/T-cell_and_glial_pathology_in_PD.

### Extraction of Nuclei and snRNAseq Procedure

Nuclei were isolated from frozen postmortem brain slices in accordance with established protocols^45,46^. Libraries were prepared using Chromium Next GEM Single Cell 3’ Reagent Kit v3.1 (PN 120237), with Chromium Single Cell A Chip Kit, 48 runs (PN 120236). Target cell recovery was 10,000 cells per sample for cingulate samples and 20,000 cells for SN samples. The final number of nuclei was calculated from the average of three counts on a Countess II or III (ThermoFisher©) using DAPI as a nuclear marker. The index plate used was 10X Dual Index Kit TT Set A (PN 1000215). Chromium Next GEM Single Cell 3LJ Reagent Kit v3.1 user guide (CG000315 Rev C) was followed. We used 10X Chromium v2 chemistry.

### Sequencing and raw data analysis

Sequencing of the resultant libraries was performed on an Illumina NOVAseq 6000 platform V4, 150 bp paired end reads, 150 cycles. Read alignment was performed using the CellRanger pipeline (v6.1.2−10X genomics) to reference GRCh38.p12 (refdata-cellranger-GRCh38-1.2.0 file provided by 10x Genomics). Count matrices were generated from BAM files using default parameters of the CellRanger pipeline^47^. Filtering and QC was performed using DecontX^48^, with default parameters, for the cingulate dataset, and CellBender^49^ for the SN dataset. CellBender (version 0.2.0) was run to remove ambient RNA with the addition of the ‘-cuda’ flag to expedite the processing. Parameters were set with an expected cell count of 10,000, total droplets included at 30,000, FPR (false-positive rate) at 0.01, and a learning rate of 0.0001, utilizing 150 epochs.

The total runtime for each sample ranged from 30 minutes to 1 hour, with acceleration achieved through the use of the NVIDIA A5000 GPU. Decontamination of background was not necessary in cingulate samples. Nuclei with percent reads aligning to mitochondrial genes >14% were excluded. Genes were filtered by keeping features with >500 counts per row in at least 100 cells. Doublets were identified using scDblFinder^50^ and then removed.

### Pre-clustering and clustering and classification of nuclei

Preclustering of nuclei was performed using Seurat’s shared nearest neighbor smart local moving algorithm^51^. First, data was normalized using SCTransform^52,53^, regressing out percent mitochondrial genes and donor. Data integration across donors was achieved using the Harmony^54^ package which effectively regressed out donor effects. Harmony embeddings were used in the FindNeighbors step. Elbow plots based on PCA for each data set were used to determine optimal number of principal components, and the Clustree package^55^ was used to determine optimal resolution values for the FindClusters() step. Seurat’s^51^ FindAllMarkers() function was used to determine basic cluster markers, which were then used to assign broad lineage identities to each cluster (astrocyte, neuron, oligodendrocyte, OPC, myeloid, endothelial, vascular, T-cell). To assist with cell type sublineage assignment, we employed EnrichR^56^, enabling us to garner information from multiple databases based on our representative genes.

Cingulate cortex neurons were assigned in line with Paryani et. al 2023^38^. Nuclei that did not conform to cell types were presumed to be doublets or artifactual noise and removed. The entire process was iteratively repeated for each lineage to remove aberrant cells and to assign subclusters, or sublineages/subtypes, within each lineage/cell type.

### Differential gene expression analysis

To compare differences in gene expression between PD and control for each cell type, we used limma^57^ within each lineage cluster. We controlled for donor, age, and sex in the model formula. Thresholds for most lineages were counts greater than 4 in at least 6 cells, and for lineages containing less than 1,000 cells, the threshold was lowered to counts greater than 2 in at least 5 cells. Our dataset did not include any separate batches. Only genes with p-values < 0.05 were carried through to downstream analyses.

### Gene set enrichment analysis and gene ontology analyses

Packages fgsea^58^ and PathfindR^59^ were used to determine gene sets enriched within our differentially expressed genes for each cell type. All differentially expressed genes along with their logFC and adjusted p values were used as input in the run_pathfindR function, using the KEGG genesets. Parameters specified were 0.05 as the adjusted p value threshold (using the adjusted p value output from limma DGE analysis), minimum gene set size 5, and maximum gene set size 500. The cluster_enriched_terms function was run, with default parameters, to find representative pathways and filter out irrelevant/uninformative pathways. Using the fgsea package, we compared our T-cell lineage to a CD8+ memory effector gene set (**Table S6**). We compiled this geneset using marker genes from the literature^60–63^. All genes in the T-cell sequencing object were assigned a logFC value through Seurat’s FindMarkers function, using PD as ident.1 and Control as ident.2, with parameters logfc.threshold, min.pct, and min.diff.pct set to -Inf to prevent filtering/removal of any genes. These genes, ranked by logFC, were input into the fgsea function with default parameters. Normalized enrichment scores and p values were reported.

To construct upset plots, we used the UpSetR^64^ package. All myeloid DEG data frames from limma voom were separated into increased (logFC>0) and decreased (logFC<0), and lists of increased and decreased DEGs were input separately into the fromList function before running the upset function with default parameters.

### Hierarchical Poisson factorization

We used the scHPF package^65^ for Python to determine interpretable factors within our SN snRNAseq dataset. The scHPF command line workflow comprises three fundamental stages: “scHPF prep,” “scHPF train,” and “scHPF score.” In the “scHPF prep” phase, the molecular count matrix is utilized to generate a matrix market file and a gene list text file. The parameter “-m” was set to 10, filtering genes to include only those present in 10 or more cells. In the “scHPF train” stage, our SN dataset was aligned with each cell type, employing a candidate parameter range from K = 7 to 17 with a step of 2. Subsequently, for the extraction of disease factors within each cell type, the training was conducted with K values of 3, 5, 7, 9, and 11. Finally, in the “scHPF score” phase, the trained models for each K value were employed to assign gene scores to individual factors, resulting in the generation of ranked gene lists. We then selected K to prevent significant overlap in gene signatures among factors. This was mainly done by observing the factors expressed by each cell type, and the K value lending itself to the most interpretable factors (gene sets following canonical gene expression patterns).

### Pseudotime analysis

Determination of single cell trajectories was performed using the Slingshot^66^ package in R. First, differentially expressed genes between fibrous-like and protoplasmic astrocytes were determined using Seurat’s FindAllMarkers function on the fibrous-like lineage, and subsetting the list of genes to those with an average log2FC greater than 0.2 (only positive log2FC values were accepted). We first ran Potential of Heat-diffusion for Affinity-based Trajectory Embedding, or PHATE^67^, dimensionality reduction on the SN astrocytes to construct PHATE embeddings using the aforementioned DEGs.

We added the PHATE embeddings to our original object and ran the slingshot function. As parameters for slingshot, we used the sublineage (protoplasmic and fibrous-like assignments) meta data for the cluster labels, and the PHATE embeddings as the dimensional reduction. All other arguments were used with their default parameters. Ridge plots were then constructed using the “slingPseudotime_1” output column, as well as the preexisting sublineage and condition meta data columns in the original object.

### Spatial transcriptomics

Following 10x Visium Spatial Protocols – Tissue Preparation Guide (CG000240), OCT embedded tissue was scored to the size of the capture area targeting the SN. One 10 µm section was mounted on each capture area of the Visium slide. Tissues on the slides were fixed using a methanol-containing buffer as per the 10X Visium manual, stained with H&E or antibodies NeuN, GFAP, and DAPI (see **Table S10** for antibody description) as per the 10X protocol for Immunofluorescence Staining & Imaging for Visium Spatial Protocols (CG000312), and then imaged. Imaging of whole slides was done at 20X magnification on a Leica DMI8 Thunder microscope. After imaging, the slides were de-cover-slipped and the tissue was permeabilized for 11LJminutes (which was empirically determined to yield best results based on the Visium Spatial Tissue Optimization Slide & Reagent Kit PN-1000193, as detailed in the protocol provided in document CG000238_RevD available in 10X demonstrated protocols). The remaining steps were conducted according to the manufacturer’s protocol to prepare the libraries. Briefly, libraries were prepared using Visium Spatial Gene Expression Slide & Reagent Kit, 16 reactions (PN-1000184). Visium Spatial Gene Expression Reagent Kits user guide (CG000239 Rev G) was followed. The libraries were sequenced on NOVAseq (paired end dual-indexed sequencing), targeting a minimum of 50,000 reads per spot.

The spatial transcriptomics (ST) samples were prepared using 10X Genomics Space Ranger (version 2.1.0) count commands, accompanied by Hematoxylin & Eosin (H&E) images in TIF format and a manually-aligned JSON file, generated from Loupe Browser (v7.0) with raw TIF images of the tissue. The loupe alignment JSON file was inputted into the loupe-alignment argument in Space Ranger along with its respective TIF image file, FASTQ reads, and slide numbers. The reference genome used for alignment was built using the Space Ranger function spaceranger mkgtf with GRCh38 as the assembly and Ensemble 91 for the transcript annotations. All other parameters were used with default settings.

### ST object preprocessing and quality control

The number of counts per spot per ST sample is shown in **Fig. S5a-j**. The plots of ST experiments shown in **Fig. 6a**, **S5** and **S6** were generated using Seurat’s SpatialFeaturePlot and SpatialDimPlot functions. A total of 10 samples were analyzed (**Table S1**). First, any spots with zero counts were removed and spot-level gene expression was normalized using SCTransform in Seurat.

### Cell type deconvolution

Deconvolution using RCTD^68^ was used to determine the proportion of each defined cell type in each ST spot from our data. As a reference, we used the normalized counts matrix and nUMI from our SN snRNAseq object with annotated cell lineages and sublineages. Queries for RCTD were generated using coordinates from the “image” and “row” columns in the Seurat object, normalized counts, and nUMI for each sample. The function run.RCTD was run with parameter doublet_mode=“full”. Otherwise, default parameters were used.

### Spatial cross-correlation

To determine how different cell types were correlated with one another on a spatial plane, we implemented spatial cross-correlation analyses^46,69^. For these analyses, we first created adjacency matrices for each sample using the getSpatialNeighbors from the MERINGUE package^70^ to denote which spots were neighbors.

To avoid false neighbor assignment of nearby cells that were not true neighbors (e.g., separated by a break in the tissue), adjacency matrices were first created using all spots, whether in tissue or not, as listed in the Space Ranger “tissue_positions” csv output file. Next, all spots assigned as “in_tissue” were kept for downstream processing, and the rest were removed. This way, spots that were not directly next to each other would not risk being labelled as first-order neighbors.

RCTD cell-type enrichment values per spot, along with each sample’s corresponding adjacency matrix, were combined to create spatial cross-correlation metrics by matrix multiplication. We used the same principles employed by MERINGUE’s^70^ spatial cross correlation function, however, due to the large sizes of our input matrices, spatial cross-correlation was implemented by matrix multiplication in Tensorflow^71^ to expedite the processing time. Specifically, the local measurement of spatial cross-correlation involves multiplying two large matrices and obtaining the diagonal elements of the resulting matrix. The speed was further enhanced by utilizing the Einsum function in the TensorFlow package, which allows for element-wise computation. The code is available at: https://github.com/dalhoomist/T-cell_and_glial_pathology_in_PD.

### Spatial transcriptomics clustering

To assign spatial clusters, we employed the R package BayesSpace^72^. We first processed our data with the spatialPreprocess function, using 7 principal components and 2000 highly variable genes for PCA, with log.normalize set to TRUE. We used the qTune function, evaluating q values between 2 and 10, and assessed the subsequent qPlot to determine the optimal number of clusters, q, defined by the elbow plot inflection point (**Table S1**). We then used the spatialCluster function on the SCT counts for each sample, using the top 7 principal components, error model t, and 1000 MCMC iterations with 100 MCMC iterations excluded as a burn-in period. All other parameters were used in their defaults.

Additionally, we sought to further classify each region of the “Surrounding_Tissue,” or all non-nigral area. We first merged all ST objects together using scCustomize and normalized them using scTransform, all as previously described in our snRNAseq processing methods. We extracted all 2000 variable features from the SCT assay. We then returned to our original objects (split by sample), and derived the spot-level gene expression values for each of the previously defined variable features. We then ran a correlation test using the “cor” function of the stats package, and generated a heatmap using the pheatmap^73^ package in R, using Manhattan distances and the ward.D clustering method (**Fig. S6k**).

### Gene set enrichment analysis in spatial data

To determine which cell types were most correlated with the Nigra and Surrounding Tissue, we employed the previously described GSEA with the package fgsea in R. We determined DEGs for each spatial region using limma voom, as described above. By using each cell type’s DEGs (logFC>0.2; positive logFC only) as a geneset, we supplied ranked genes from each region.

We also measured the spot-level enrichment values for our T-cell disease factor. We employed the fgsea package as described above, comparing each spot’s gene expression to the top 200 ranked genes from the T-cell disease factor.

### Immunohistochemistry and histology

To validate our findings that T-cells assume a tissue resident memory (TRM) phenotype in Parkinson’s disease, we performed immunohistochemical staining for TRM-specific marker CD103^74^ in postmortem control and Parkinson’s human brain sections, 7 mm thick (for antibody description, see **Table S10**). The SN was analyzed in transverse sections of the midbrain at the level of the red nucleus. All immunostains were conducted on a Leica© Bond RXm automated stainer. For chromogenic DAB stains, a generic IHC protocol was employed as per manufacturer protocols. Standard deparaffinization and rehydration steps preceded antigen retrieval in Leica ER2 (Cat. No. AR9640) antigen retrieval buffer for 10-20 minutes according to manufacturer protocols. Then, a peroxide block was applied for 10 minutes followed by three wash steps using bond wash solution (Cat. No. AR9590). A one-hour incubation in a blocking buffer in 10% donkey serum containing PBS-based buffer preceded antibody labeling for 1 hour at ambient temperature. This was followed by three wash steps after which the Post Primary was dispersed for 8 minutes, followed by three wash steps prior to the Polymer being dispersed for 8 minutes, followed by another three wash steps. The slides were then treated with deionized water for one minute prior to incubating in Mixed DAB refined for 10 minutes followed by three washes of deionized water. Slides were stained with Hematoxylin for five minutes followed by a wash with deionized water, then Bond wash solution and a lastly deionized water wash. For multiplexing immunostains using antibodies raised in non-overlapping hosts, we used a generic immunofluorescence protocol. Slides were baked in a 65 °C oven for a minimum of 2 hours. The following protocol was then used: After a dewaxing step, incubation in BOND Epitope Retrieval Solution 2 (Cat. No. AR9640) for 20 minutes was used for heat-induced epitope retrieval. Next, the slides were washed in 1X PBS before washing twice in Bond Wash Solution (Cat. No. AR9590) – 10 minutes/wash. Next, they were incubated in 10% donkey blocking serum for 60 minutes followed by the primary antibody diluted in blocking buffer for 60 minutes. After three washes, the slides were incubated in the secondary antibody containing buffer for 60 minutes. After three washes, a DAPI containing mounting solution (Everbright TrueBlack Hardset Mounting Medium with DAPI, Cat. No. 23018) was used to label nuclei and quench autofluorescence prior to cover-slipping. A volume of 150 ml/slide was used for all steps. All steps were conducted at ambient temperature.

Brightfield images were acquired with a Leica Aperio LSM™ slide scanner under 20X objective. All immunofluorescent images were acquired on the Leica Thunder imager DMi8. Images were acquired at 20X using a Leica K5 camera. Leica Biosystems LAS X software was used for image capture. Tiles covering the cingulate and SN were taken and stitched. Leica Thunder instant computational clearing was used to remove out of focus light.

### Quantification of IHC

All image analysis was performed in QuPath 0.42^75^. Annotations detailing the cingulate, peduncle or SN were manually drawn. To detect cells, we used the “cell detection” function under the analysis menu, with DAPI as the Detection Channel. We modified the background threshold per image to eliminate non-specific detections. We then trained an object classifier to classify the detections for the different channels. Training data was created from each image to delineate cells that were positive for the specific antigens in question. One classifier per channel was trained by calling the “train object classifier” function with the following parameters: type = Random Trees, measurements = Cell: Channel X standard deviation, mean, max, and min measurements for the channel in question. To increase the accuracy of the classifier, additional training annotations were created on the image in question until the classification results matched the impression of the observer. Once a classifier was trained for each channel, “create composite classifier” was called to create a classifier consisting of multiple individual classifiers, one for each channel on the image. Classifiers were trained for each image separately. For the DAB stains, positive cell detection was used by detecting optical density sum to detect nuclei for CD8+ cells. An object classifier was again trained by using the “train object classifier” function, with the following parameters: type=Random Trees, measurements = all measurements, and selected classes= CD8+ and CD8-. The number of cells identified as CD8+ were then normalized by dividing by the area of the annotation in which the analysis was done. CD103 quantification was also performed. The numbers of CD103+ cells in the SN were counted manually by two board certified neuropathologists (OAD, JEG). These counts were then divided by the area of the respective region. All statistical analyses were conducted in GraphPad® Prism 10. One-tailed or two-tailed unpaired t-tests were used to compare PD vs control (Fig. 3, Fig. S7), as indicated in the figure legends. One-tailed t-test was used when we had a prior hypothesis informed by the transcriptomic data.

### Statistical testing

Statistical testing, aside from IHC analyses above, were conducted using R version 4.2.2. Linear model regression analyses seen in **Fig. 1c**, **Fig. 6c**, and **Fig. S1e** were conducted using the lm function of the stats package. Prior to linear model testing, data were tested for normality using the shapiro.test function from the stats package with default parameters. Wilcox testing seen in **Fig. 1f** was conducted using the wilcox.test function in the stats package with default parameters. T tests, as seen in **Fig. S1c-e**, used the t.test function from the stats package to run a Welch Two Sample t-test, with the alternative hypothesis being that the true difference in mean between group Control and group PD is less than 0 for **Fig. S1c**, and two-sided for **S1d-e**. Asymptotic two-sample Kolmogorov-Smirnov seen in **Fig. 2d** and **2h** were run using default parameters in ks.test function from stats package. Statistics comparing PD to control for **Fig. 3** and **Fig. S7** were done using GraphPad® Prism 10 using one-tailed unpaired t-tests.

**Figure 1.**
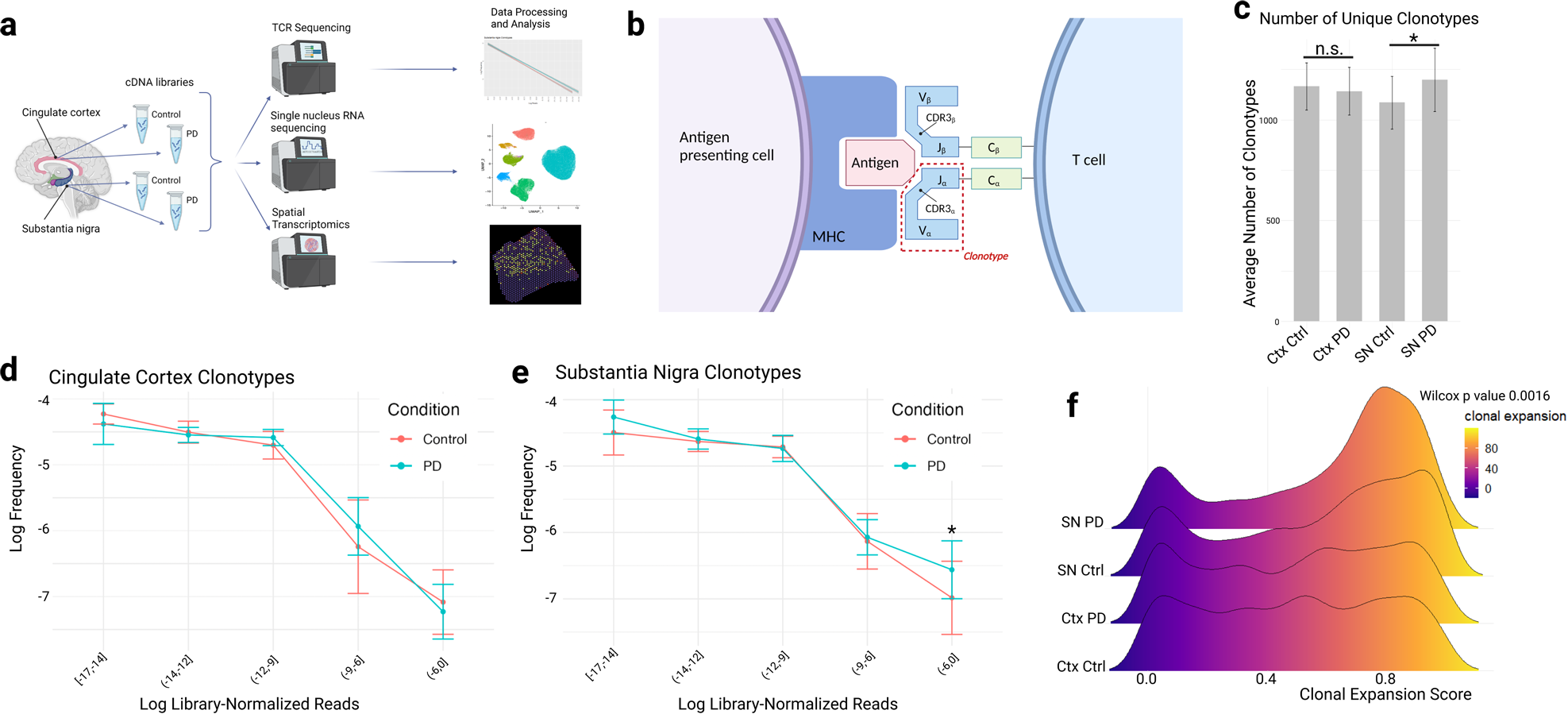
T-cell receptor sequencing identifies clonal expansion in T-cells of the substantia nigra in Parkinson’s. **a)** Schematic of the study design. Cingulate cortex and substantia nigra (SN) tissue samples were dissected from PD and control frozen postmortem brains, and then processed by TCR sequencing, snRNAseq, and spatial transcriptomics, and analyzed computationally. **b)** Diagram of a T-cell receptor. The alpha chain is shown, where V is the variable region, J the joining region, C the constant region, and CDR3 (complementarity-determining-3 region). An antigen is presented by MHC-II complex and recognized by the T-cell receptor on the right. A clonotype is defined as the combination of the V, J, and CDR3 regions. **c)** Bar graph depicting the number of unique clonotypes detected in the cingulate cortex and the SN, PD and control. The difference between conditions in the cingulate is not significant (linear regression coefficient estimate – 24.83 and p value 0.689) but is significant in the SN (linear regression coefficient estimate 112.75 and p value 0.0479). **d-e)** Power law graphs of cingulate cortex and SN clonotypes, respectively, with bins of the log number of library-normalized reads on the x axis, and log of frequency (library-normalized) of clonotypes for each bin expressed on the y axis. Control clonotypes are represented in orange, and PD in blue. Error bars represent standard deviation. **f)** A ridge plot depicting clonal expansion score as calculated by GLIPH2 for each region and condition. Wilcox test derived p value from comparing the average expansion score of the PD vs control distribution in the substantia nigra is 0.0016.

**Figure 2.**
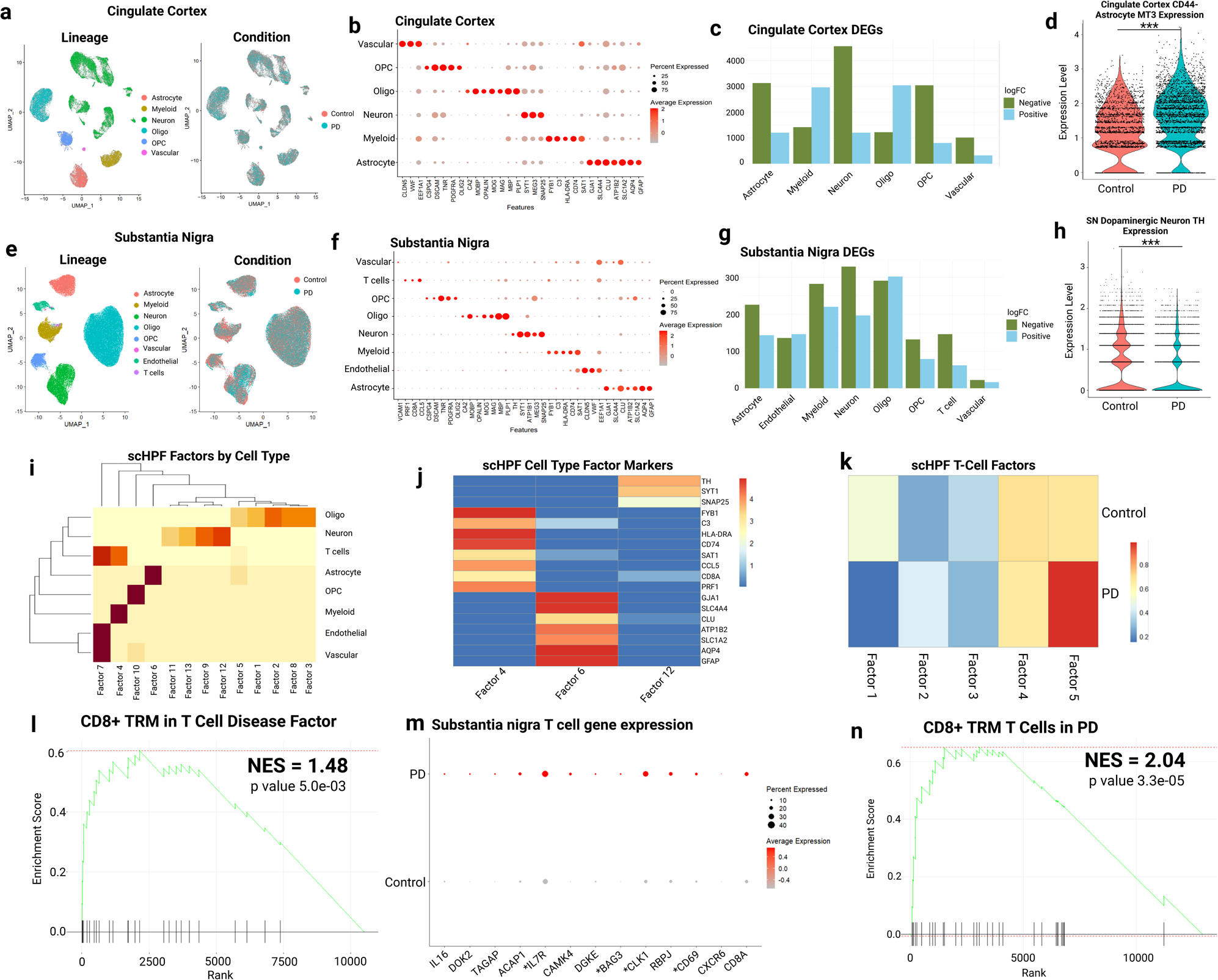
Single nucleus RNA sequencing reports differences in gene expression patterns of PD lineages and identifies a PD T-cell signature. **a)** Uniform manifold approximation and projection (UMAP) graphs showing nuclei from the cingulate cortex, PD and control, grouped by assigned lineage and by condition. **b)** Dot plot of select gene (x-axis) marker expression in major lineages in the cingulate cortex (y axis). Size indicates percentage expression, and color indicates normalized expression levels. **c)** Bar plot of the number of significant differentially expressed genes for all lineages in the cingulate cortex in PD compared to control, with downregulated genes (negative log fold change – green), and upregulated genes (positive log fold change - blue). **d)** Violin plot showing the gene expression of *MT3* in control (orange) and PD (blue) in protoplasmic astrocytes in the cingulate cortex (logFC PD vs control: −0.468, p value 3.37e-27). **e)** Same as **a**, but for SN nuclei. **f)** Same as **b** but for the SN. **g)** Same as c, but for the SN. **h)** Same as **d** but for *TH* SN dopaminergic neurons (logFC PD vs control: –0.23, p value 1.35e-28). **i)** Heatmap showing scores single cell hierarchical Poisson factorization (scHPF) gene factors (columns) projected on lineages (rows). **j)** Heatmap showing the normalized gene expression for select cell type markers in select scHPF factors. **k)** Heatmap of average cell score of PD and control nuclei in each T-cell scHPF factor. Columns represent factors, rows represent condition. Blue indicates a low score, red indicates a high enrichment score. **l)** Gene set enrichment analysis of the *CD8*+ T-cell resident memory (TRM) gene set in T-cell disease factor (factor 5), with normalized enrichment score (NES) of 1.48 and p value 5.0e-03. **m)** Dot plot of genes expressed in PD and control T-cells. Asterisks (*) denote genes differentially expressed in PD. **n)** Pre-ranked gene set enrichment analysis of the *CD8*+ TRM gene set in the T-cell gene expression ranked by logFC in PD vs control. P value 3.3-e05, and normalized enrichment score (NES) 2.04.

To determine modality as seen in **Fig. S1i**, the function Modes from LaplacesDemon^76^ R package was used on each distribution of clonal expansion scores. Further, we ran an asymptotic two-sample Kolmogorov-Smirnov test on the distributions using the ks.test function from the stats package with default parameters.

### Figure Generation

All figures were created with Biorender.com, or GIMP version 2.10.

## Results

### T-cell receptor sequencing data reveals clonal expansion in the substantia nigra of Parkinson’s disease subjects

The numbers of T-cells are increased in the SN of subjects with PD^5,6^ and the cortex of Diffuse Lewy Body Disease (DLBD)^29^. To examine whether T-cells in the PD/DLBD brain display clonal expansion and/or increased clonal diversity, we compared the T-cell receptor (TCR) repertoires in PD/DLBD and control samples using transcriptomics and TCR sequencing (Fig. 1a) in 44 brain donors.

We compared TCR repertoires in the SN to those in the cingulate cortex, a region commonly containing Lewy bodies in advanced PD with dementia (PDD) and DLBD. We chose the cingulate cortex in PDD/DLBD because it exhibits neurodegeneration, but to our knowledge has not been reported to display increased T-cells in these disorders. We analyzed a total of 44 samples from 44 patients from either cingulate cortex or SN (Cingulate: n=10 PD and 6 controls, SN: n=13 PD and 15 controls - see **Table S1** for demographic data). Note, we did not have paired SN and cingulate samples in our cohort, and some controls had other neuropathological changes that did not include a synucleinopathy. For simplicity, we refer to cortical PDD/DLBD as PD thenceforward.

Alpha and beta chains of TCRs are highly correlated^39^, and here we sequenced the TCR alpha chain (Fig. 1b). As a quality control step, we determined that all libraries were fully saturated, with adequate read depths and numbers of sequences (**Table S2**). We first compared the number of unique clonotype repertoires in PD and control SN and cingulate using a linear model. There was a significantly higher number of unique clonotypes in PD than control in the SN, but no significant difference in the cingulate (Fig. 1c). We were also interested in the CDR3 sequences that were shared amongst individuals (public, or here referred to as global sequences)^77^. We identified some overlap among brain regions and conditions in these global CDR3 sequences (**Fig. S1a**), however most global CDR3s were of low abundance (**Fig. S1b**). Still, SN PD showed more global CDR3 sequences compared to controls (**Fig. S1c**).

We next assessed Shannon entropy levels in our repertoires which, through calculating the probability of each CDR3 sequence, determined the diversity of each repertoire^78^ (see methods and supplementary results), and we found increased entropy in the SN compared with the cingulate (**Fig. S1d-e**). Together, these data all indicate that there are more TCR sequences in the PD SN than compared with the cingulate cortex.

To determine if T-cells show features of clonal expansion in PD, we plotted the frequency of TCR clonotypes at a given read depth against the read depth (Fig. 1d**-e**). As expected, most clonotypes were of low read depth, which is consistent with previous work^77–79^. Interestingly, we found a significant increase in the proportion of most abundant clonotypes (defined as log10 normalized read depth greater than −6) in PD SN compared to control SN. This was not observed in the PD cingulate cortex. Together, these data indicate that PD subjects possess more clonotypes that are relatively more abundant in the SN, consistent with clonal expansion.

As another measure of clonal expansion, we devised a measure that leverages the global and local distributions of abundance of each repertoire. The patient-level clonal score essentially compares the deviation of the observed clonotype abundance from a hypothetical random distribution derived from the entire dataset (see methods). For each patient, we compared the deviation of the repertoire distribution from a random distribution using a Kolmogorov-Smirnov test, and then compared the resulting D-values (clonal expansion scores) across conditions and regions. We found increased clonal expansion in PD versus control SN but not in the cingulate cortex (**Fig. S1f**). Further, in assessing the p value distribution from 100 repetitions of testing, the data showed a consistent trend of significance in the SN, and insignificance in the cingulate (**Fig. S1g**). Together, these data further support that T-cells show features of clonal expansion in the PD SN.

In the periphery, T-cells that recognize alpha-synuclein have a broad diversity of TCRs and no public clones^7^. As blood samples were not acquired from the subjects prior to death, we could not directly identify which peripheral TCR sequences had been expanded in the brain. To analyze the similarity amongst TCRs in our dataset, we employed GLIPH2^44^ analysis to cluster CDR3s based on amino acid feature homology. This analysis generates clusters/motifs of CDR3 sequences that can have members derived from different samples and assigns each motif a clonal expansion score. We identified 16,875 global motifs, i.e., strings of conserved amino acids with a single amino acid substitution, and 3,103 local clusters, i.e., uninterrupted sequences of amino acids.

We found extensive diversity in the sizes of the clusters, ranging from the minimum of two CDR3 sequences to 4,698 copies with a motif, with a median value of two CDR3 sequences. (**Table S3**). We filtered our results for significance (Fisher score < 0.05), and determined a representative condition for each tag, or conserved motif, depending on which condition contributed the highest proportion of CDR3 sequences in the tag. We found that motifs predominated by SN PD CDR3 were more likely to exhibit high clonal expansion scores compared to control SN predominated motifs (Wilcox p value 0.0016; Fig. 1f).

Taken together, this analysis revealed a significant increase in the patient-level clonal expansion score in PD SN (**Fig. S1f-g**), indicating T-cells are clonally expanded in the PD SN.

### Single nucleus RNA sequencing reveals cell-type specific DEGs in PD

We then examined the T-cell gene signatures with single nucleus RNA sequencing (snRNAseq) (Cingulate: n=10 PD and 8 controls, SN: n=13 PD and 15 controls -**Table S1)**. The SN dataset includes 207,859 nuclei, with 96,244 derived from PD subjects (including 831 SN T-cells, 535 being from PD donors), and the cingulate dataset comprises 57,425 nuclei, 32,442 from PD subjects. We projected these nuclei in UMAP space and assigned cell types/lineages in the cingulate and SN (Fig. 2a for the cingulate, and Fig. 2e for the SN), and by donor and sex (**Fig. S2a** for the SN, and **Fig. S2e** for the cingulate). The expression of select canonical marker genes per lineage for the cingulate and SN is shown Fig. 2b and Fig. 2f, respectively, and cluster markers are reported in **Tables S4-5** for the SN and cingulate, respectively.

We determined the differentially expressed genes (DEGs) between PD and control in each lineage in the SN and cingulate (**Table S4** and **Table S5**). We found that the largest number of altered DEGs in PD were in neurons and oligodendrocytes in both SN and the cingulate (**Fig.2c** and Fig. 2g). There were also high numbers of DEGs in astrocytes and myeloid cells in both regions, and in T-cells in the SN. Interestingly, astrocytes showed increased *MT3* expression in the cingulate in PD (Fig. 2d), as we have shown for Huntington’s disease (HD)^38^. We validated this finding in our supplemental IHC data (**Fig. S7g**). As expected, SN neurons exhibited reduced expression of the dopamine synthetic enzyme tyrosine hydroxylase (*TH*) (Fig. 2h), consistent with previous studies^3^.

Prior studies have detailed neuronal and glial pathology in PD at the single nucleus level^33,36^, including recent preprints^34,35,80^. We described our sequencing results from neurons in the SN (**Fig. S2a-d**) and the cingulate cortex (**Fig. S2e-h**), and the pathology associated with them (**Fig. S3a-d**) in the supplementary results. Briefly, we found that most DEGs were in SN TH+ neurons in the SN and cingulate layer 2-3 CUX2+ glutamatergic neurons (**Fig. S2d** and **Fig. S2h**, respectively), and several pathways involved in neurodegeneration and synaptic vesicle cycle that were shared between the two most affected neuronal populations (**Fig. S3a-d**).

### Single nucleus RNA sequencing defines a T-cell PD disease signature and CD8+ resident memory phenotype

We analyzed our dataset using a recently developed approach called single cell Hierarchical Poisson Factorization^65^ (scHPF; see methods). This method derives factors, or gene sets, that capture the sources of gene expression variability in the dataset, which could be lineage related, disease related, or related to other factors. When we applied scHPF to the SN snRNAseq dataset, we retrieved factors that corresponded to cell types (Fig. 2i). An example of the gene score of select factors is shown in Fig. 2j, where genes from astrocytic, TH+ neuron, and T-cell factors are shown, underscoring the power and validity of the technique. Additionally, T-cells in our dataset were mostly CD8+ (Fig. 2f), consistent with previous reports^81,82^ and our validation studies (see below; **Fig. 3a-d**).

**Figure 3:**
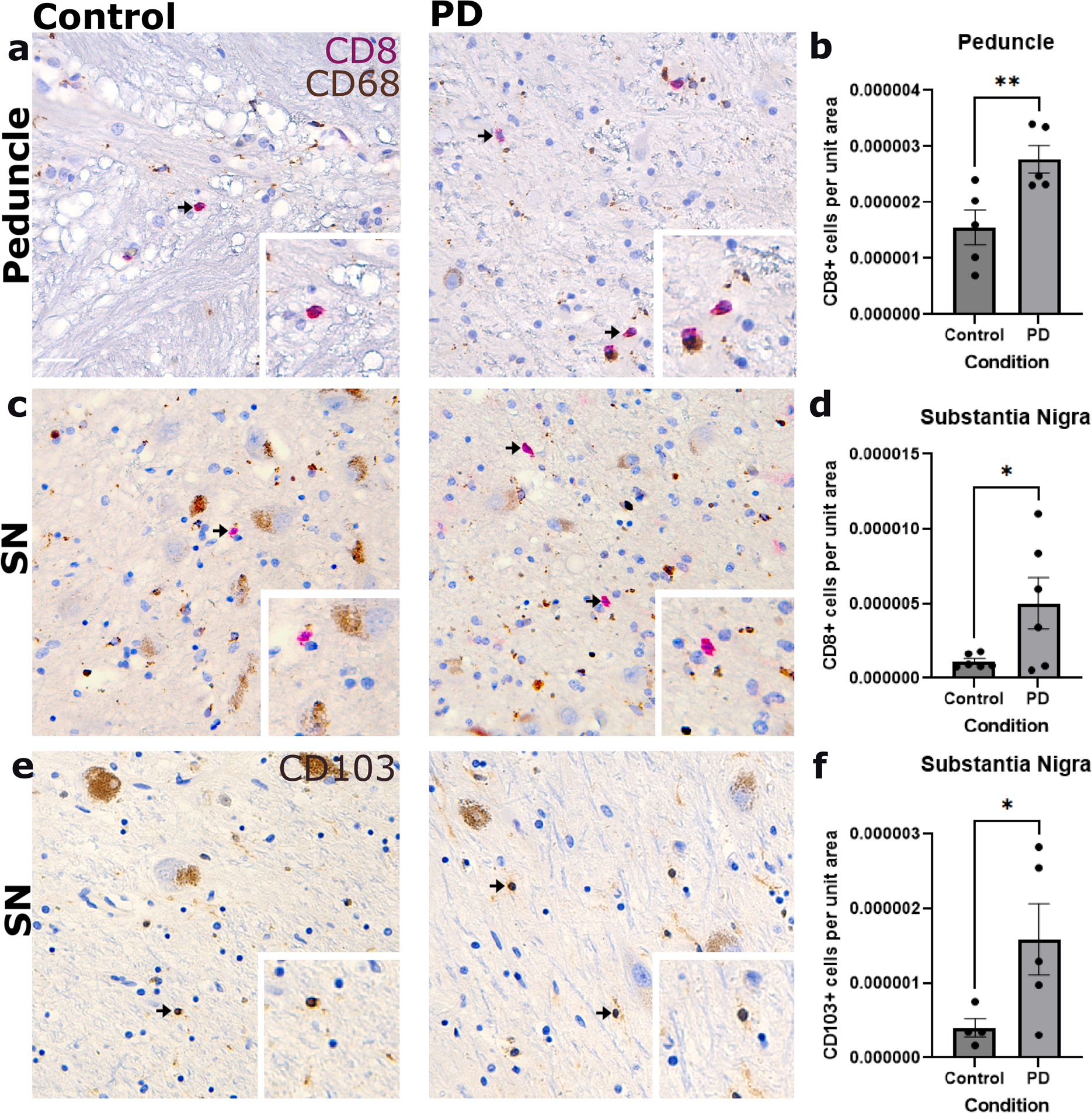
Validation of T-cell phenotypes in the SN. **a)** Immunohistochemical stains for CD8 (red) and CD68 (brown) in the cerebral peduncle. Scale: 50 μm. **b)** Quantification of the density of CD8 positive cells per unit area in the peduncle. Unpaired one-tailed T-test with N=6 for control and PD. P value = 0.0079. Data is shown as mean +/−SEM. **c)** Immunohistochemical stains for CD8 (red) in the SN. **d)** Quantification of the density of CD8 positive cells per unit area in the SN. Unpaired one-tailed T-test with N = 6 for Control and PD. P value = 0.0239. Data is shown as mean +/−SEM. **e)** Immunohistochemical stains for CD103 (brown) in the SN. **f)** Quantification of the density of CD103 positive cells per unit area in the SN. Unpaired one-tailed T-test with N = 4 for control and N = 5 for PD. P value = 0.0343. Data is shown as mean +/−SEM.

We then used scHPF to extract a PD “disease factor” (**Table S7**) in T-cells with higher scores for PD than control subjects (Fig. 2k). Factor 1 was higher in control T-cells, while PD T-cells had higher scores in factor 5, nominating this factor as a “disease factor”. The top genes in factor 5 are implicated in IL-2 signaling (*UBC, SOS1, CD2, JAK3, LCK, BIRC3, DOK2, HIST1H2AC*) and NFkB signaling (*TNFAIP3, RIPK1, CFLAR, PLCG2, LCK*). Given the known effects of IL-2 signaling on CD8+ T-cell fates, including effector and memory phenotypes^83^, we performed GSEA analysis to measure enrichment of a T-cell resident memory (TRM) gene set^63,84–93^ (**Table S6**; Fig. 2l) in the T-cell disease factor 5. We found that the gene set was enriched in the T-cell disease factor. Also, the expression of multiple genes from the TRM gene set were significantly increased in PD T-cells, including T-cell memory genes such as *IL7R* and *CD69*^88,94^ (Fig. 2m).

As expected, the TRM gene set was significantly enriched in the gene expression of PD T-cells ranked by the log-fold change (logFC) from control T-cells (Fig. 2n). We also assessed the enrichment of a general memory T-cell gene set (**Table S6)**, encompassing central memory, effector memory, and peripheral memory T-cell genes^95–97^ in T-cells genes ranked by their gene-wise logFC in PD vs control, and a found a similar enrichment in PD **(Fig. S1j**), further corroborating our findings that T-cells demonstrate a TRM phenotype in the PD SN.

Taken together, these results indicate that PD SN T-cells demonstrate a more prominent memory phenotype, which we interpret as being more antigen-experienced.

### Validation of T-cell phenotypes in the post-mortem substantia nigra

To independently validate that PD showed increased T-cells are memory CD8*+* cells, we performed IHC on post-mortem SN tissue. We labeled cytotoxic T-cells with a CD8 antibody and found that the density of CD8+ T-cells was higher in the SN parenchyma and in the white matter (cerebral peduncle) in PD brains compared to controls (Fig. 3a**-d**), consistent with previous reports showing that CD8+ T-cells infiltrate the PD brain^5,6^. The cerebral peduncle is useful in these analyses, as blood vessels enter the SN from the subarachnoid space around the midbrain via the peduncle, and many CD8+ T-cells can be found around these vessels in the peduncle and peduncular tissue itself. We did not count cells around vessels, thus, the T-cells we quantified in the SN and in the white matter were parenchymal.

To validate the memory phenotype of CD8+ T-cells in PD SN, we used an antibody against CD103, which is expressed in TRMs^98^. The density of CD103*+* cells was higher in PD compared to controls in the SN (**Fig. 3e-f**), supporting the transcriptomic results indicating that T-cells in the PD SN adopt a memory resident phenotype.

### Myeloid cells in the PD SN show increased enrichment in neuroinflammatory pathways

We then analyzed SN myeloid/immune cells in isolation through subclustering and DGE analysis. We identified three myeloid states: baseline/homeostatic myeloid cells, activated microglia/myeloid cells, and monocyte-like myeloid cells/border-associated macrophages (BAMs) (**Fig. 4a**). Select markers of each subcluster are shown in **Fig. 4b** (**Table S4**). We found that activated microglia/myeloid cells exhibited the highest number of DEGs in PD, and that baseline myeloid cells and monocytes/BAMs demonstrated lower numbers of DEGs (Fig 4c; **Table S4**). A recent preprint identified heterogeneous transcriptional states in the PD SN through higher resolution clustering^34^. For this study, we find it more expedient to compare the broad classes of myeloid cells highlighted herein.

**Figure 4.**
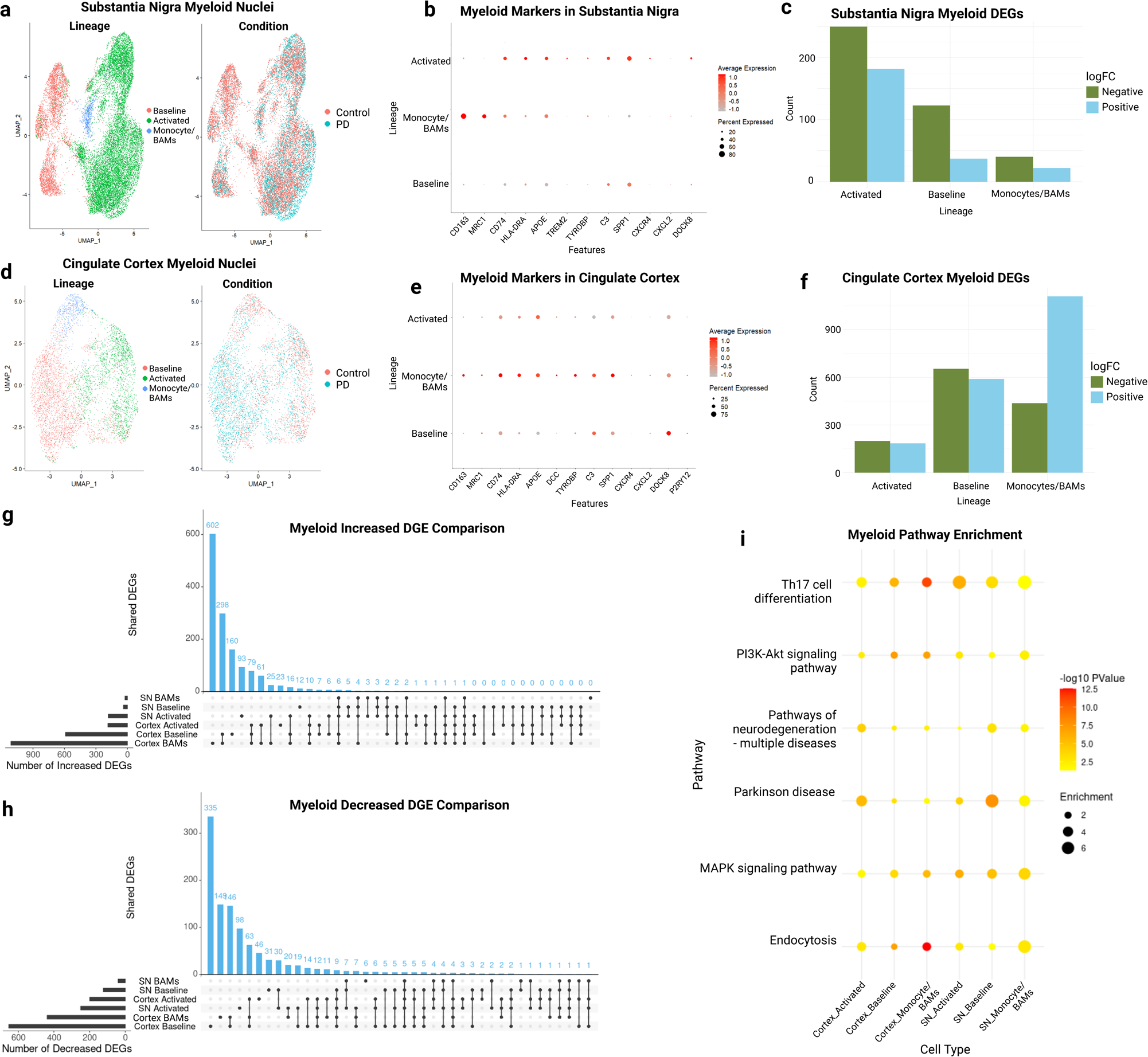
Patterns of dysregulation of myeloid cells in the substantia nigra and cingulate cortex. **a)** UMAP plots of myeloid cells in the substantia nigra (SN) from snRNAseq, grouped by lineage/sublineage (left), and condition (right). **b)** Dot plot of select gene (x-axis) marker expression in myeloid lineages in the SN (y axis). Size indicates percentage expression, and color indicates normalized expression levels. **c)** Bar plot depicting number of differentially expressed genes in the PD SN on the y axis and each myeloid lineage on the x axis. Green bars represent DEGs with a negative logFC, or decreased expression in PD, and blue bars represent DEGs with a positive logFC, or increased expression in PD. **d)** Same as **a**, but for cingulate cortex myeloid cells. **e)** Same as **b**, but for cingulate cortex. **f)** Same as c, but for cingulate cortex. **g-h)** Upset plot showing the patterns of overlap between DEGs increased (**g**) and decreased (**h**) in PD across different myeloid sublineages (rows) in the SN and cingulate cortex. Number of shared DEGs are plotted in blue along the y-axis – note that BAMs refers to monocytes/BAMs and has been shortened for improved visualization. The cell type combinations between which the DEGs are shared are displayed across the x axis, with black dots representing cell types present in the combination, and light gray dots representing cell types not present in the combination. Number of total increased DEGs are plotted to the left of each cell type name. **i)** Dot plot showing KEGG pathway enrichment scores and adjusted p values of select pathways of each myeloid sublineage in the SN and cingulate cortex. Pathways are shown on the y axis term names, and the x axis shows each myeloid sublineage. The size of each dot represents its fold enrichment value, and the color represents its –log10 p value, with yellow denoting lower significance and red indicating higher significance.

To determine how SN myeloid cells are affected in PD, we compared activated, baseline, and monocyte/BAM subtypes to the myeloid cells of the cingulate (**Fig. 4d-e**; **Table S5**). The cingulate cortex demonstrated much higher numbers of DEGs across all myeloid lineages. Relatively, the activated myeloid cells had the highest number of DEGs in the SN, and baseline myeloid cells and monocytes/BAMs had the highest number of DEGs in the cingulate (**Fig. 4f**; all cingulate DEGs **Table S5**).

To compare the response of myeloid cells in the cingulate and SN, we examined the patterns of overlap between positive and negative DEGs in Upset plots (**Fig. 4g-h**; see methods). We found that the majority of DEGs were region-specific: in the cingulate there were 142 DEGs (79 increased and 63 decreased) shared between all cortical myeloid cells and no SN myeloid cells (the DEGs, both increased and decreased, shared between myeloid lineages in the cortex and SN can be found in **Table S9**). The SN had much lower numbers of DEGs in general, with just 10 DEGs (3 increased and 7 decreased) specific to SN myeloid cells. Most other DEGs were shared across cell types or brain regions. This data points to a region-specific myeloid gene signature in the cingulate cortex in PD, but a non-specific response in the PD SN (**Fig. 4g-h**; **Table S9**).

To decipher these gene sets, we performed gene set enrichment analysis of all KEGG pathways with the pathfindR package and found several shared dysregulated pathways (**Fig. 4i**). SN monocytes/BAMs showed some of the highest enrichment scores in Th17 cell differentiation, PI3K-Akt signaling pathway, MAPK signaling pathway, and endocytosis. As PI3K-Akt and MAPK signaling are involved in neuroinflammation in neurodegeneration and are potential therapeutic targets in PD^99–101^, these data suggest that monocytes/BAMs in the SN participate in neuroinflammation and immune signaling in PD.

### Differential regional dysregulation of astrocytes in PD

Astrocytes play roles in PD^102^. We and others have shown that astrocytes can be distinguished by *CD44* expression into fibrous-like and protoplasmic^103,104^. We assigned astrocytic nuclei to a protoplasmic or fibrous-like sublineage and conducted DGE analysis (**Fig. 5a-c** for SN, and **Fig. S4a-b** for cingulate cortex). Following our previous results that indicated a compensatory neuroprotective astrocytic response characterized by increased metallothionein protein MT3 expression in HD astrocytes^38,45^, we found that *MT3* was increased in the CD44- (protoplasmic) cingulate cortex astrocytes but not SN protoplasmic astrocytes. As expected, expression of *GFAP* was increased in both regions (**Fig. 5d**).

**Figure 5.**
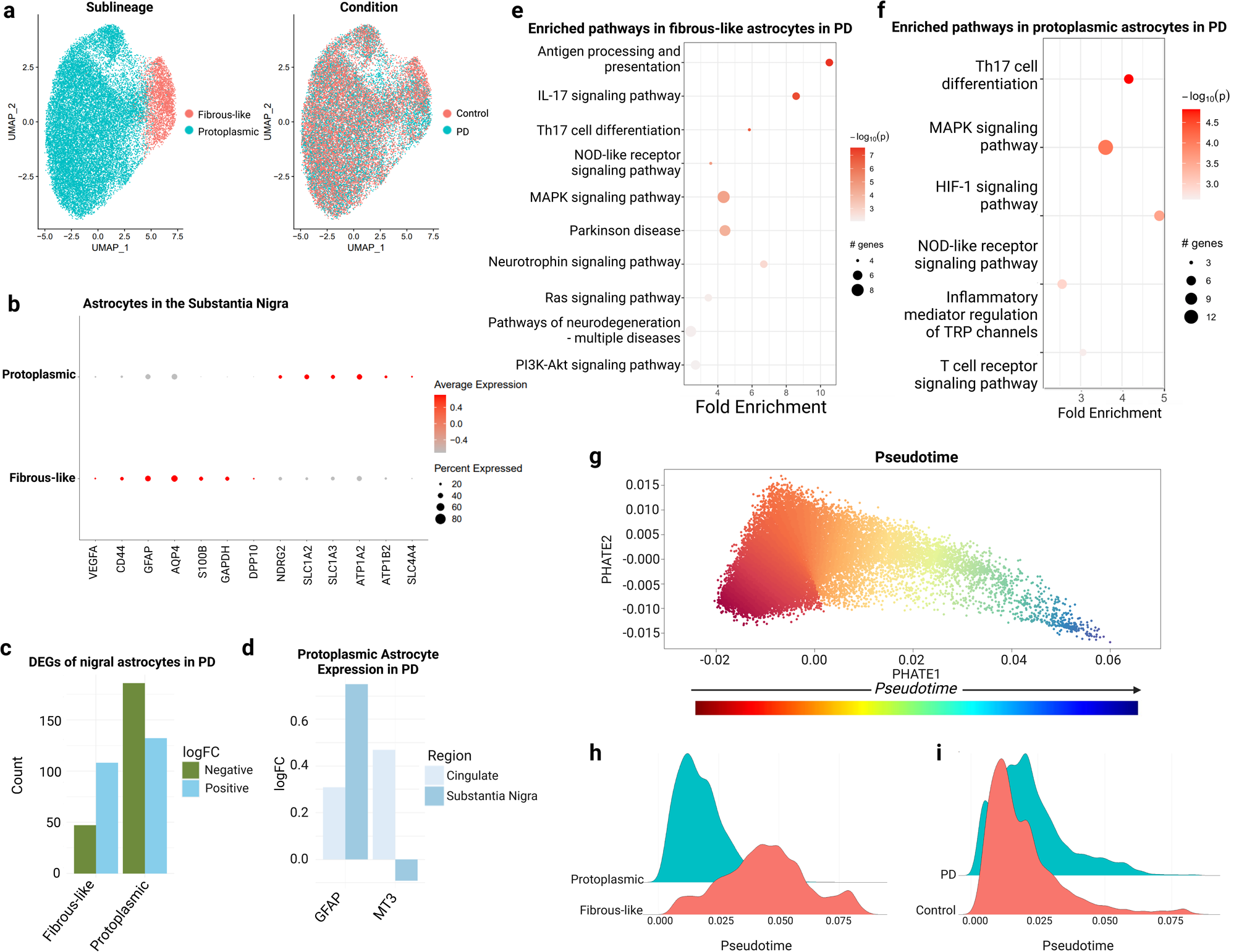
Single nucleus RNA sequencing of substantia nigra astrocytes in PD indicates abnormal functioning. **a)** UMAP plots of nigral astrocytes, grouped by sublineage (left) and condition (right). **b)** Dot plot with select markers genes along the x axis and astrocyte sublineages on the y axis. Size percent of expression, and color denotes normalized expression levels. **c)** Bar plot depicting the number of differentially expressed genes in PD for astrocyte sublineages. Green bars represent DEGs with a negative logFC, or with decreased expression in PD, and blue bars represent DEGs with a positive logFC, or increased expression in PD. Fibrous-like astrocytes are shown on the left, protoplasmic on the right. **d)** Bar plot of *GFAP* and *MT3* differential expression in PD in cingulate cortex and SN protoplasmic astrocytes. The cingulate cortex is denoted by light blue bars, the nigra by dark blue bars. *MT3* is decreased in protoplasmic astrocytes in the PD SN (logFC –0.09 and p value 1.4e-10), and increased in PD cingulate protoplasmic astrocytes (logFC 0.47 and p value 3.4e-270. GFAP is increased in both PD SN and cingulate cortex protoplasmic astrocytes (logFC 0.75 and p value 6.4e-290 - SN, logFC 0.31 and p value 7.8e-12 – cingulate cortex). **e-f)** KEGG pathway enrichment analysis of SN fibrous-like (**e**) and protoplasmic (**f**) astrocytes PD DEGs. **g)** Pseudotime plot of astrocytic nuclei in the SN, with protoplasmic astrocytes on the left, and fibrous-like astrocytes on the right. Nuclei are projected on PHATE axes. Color bar indicates pseudotime value range (red is low and blue is high). **h-i)** Ridge plot depicting the proportion of protoplasmic and fibrous-like (h) and PD vs control (i) at each pseudotime value.

We next sought to validate these findings with immunohistochemical studies in post-mortem brain specimens. To do so, we immunostained cingulate cortex and SN, PD and control, with GFAP and MT3 (**Fig. S7a-f**). We found that MT3 was significantly increased in GFAP-high astrocytes in the cingulate cortex in PD, but there was no such significant increase in the SN in PD, confirming our gene expression data. Interestingly, when we quantified the proportion of astrocytes that were GFAP-high – a surrogate for reactive astrocytes – we found that more astrocytes were GFAP-high in the cingulate, but not the SN. In fact, there was a reduction in the proportion of GFAP-high astrocytes in the SN (**Fig. S7g**). This is compatible with previous reports^105,106^ that found reduced GFAP protein expression in the PD SN.

We then examined KEGG pathways enriched in the DEGs (both increased and decreased) in the SN fibrous-like and protoplasmic astrocytes (**Fig. 5e-f**; **Table S4**) and in the cingulate cortex (**Fig S4c**; **Table S5**). In both astrocyte types and both brain regions, we found enrichment in multiple immune activation pathways including IL-17 signaling and NOD-like receptor signaling pathways in PD. These were largely driven by DEGs increased in PD including *FOS* and *JUN*, which are known to be increased by several pathways, including stimulation of astrocytes by interferon gamma, IL-1, and IL-6 amongst others^107^. Other shared pathways include MAPK signaling and pathways related to stemness (**Tables S4-5**). Of note, the latter pathway was driven by different DEGs in the cortex versus the SN. For instance, in fibrous-like cingulate astrocytes, it was driven by downregulation of *FGFR2*, *FGFR3*, and *BMP* receptor-1 genes. Conversely, in SN fibrous-like astrocytes, it was driven by upregulation of *ID3*, *ID4* and downregulation of *LIFR*. Interestingly, VEGF-signaling was enriched in both astrocyte types in the cingulate (**Table S5)** but only fibrous-like astrocytes in the SN (**Table S4**), where it was driven by upregulation of *FOS, JUN, VEGF*, and *PLCG2*, and downregulation of *TXNIP, COX2*, and *CALM2*. Further, enrichment of pathways related to neurodegeneration was higher in fibrous astrocytes in both brain regions (**Fig. 5e-f**; **Fig. S4c; Tables S4** and **S5**).

We then examined evidence for a transition from protoplasmic to fibrous-like states in PD, as we have seen in HD, hypoxia, and seizures^38,103^, using pseudotime analysis which is a way to order cells along a trajectory of gene signature. We ordered cells on the axis of variation along the genes that differ between the two astrocyte sublineages (**Fig. 5g-h** – SN, and **Fig. S4e-f** – cingulate cortex). We found a trajectory from protoplasmic to fibrous-like (**Fig. 5h** – SN, and **Fig. S4f** – cingulate cortex), with protoplasmic at the start of the trajectory, and fibrous-like at the end. At the population level, there appears to be a shift towards fibrous-like phenotypes in PD astrocytes, with Wilcox p value <2.2e-16 for astrocytes in both the SN (**Fig. 5i**) and in the cingulate cortex (**Fig. S4g**), which is consistent with a previous report^33^. However, in taking the mean or median from each donor and comparing at the patient level, we do not observe a significant increase in pseudotime values between conditions (one-sided Wilcox p value 0.4 for SN, 0.2 for cingulate). In future studies, a larger dataset may further highlight a fibrous-like transition in PD astrocytes. Further, as seen in HD, there are regional differences in astrocytic responses to injury, with the most severely affected regions showing no increase in the neuroprotective gene *MT3*.

### Spatial transcriptomics reveals spatially diverse patterns of pathology in PD

To spatially map the disease signature within the diverse brain microenvironments that harbor these cells, we conducted spatial transcriptomics on a subset of our SN tissue samples (**Fig. S5a-j**). First, to evaluate cell-type-specific gene signatures in our spatial transcriptomics data, we employed Robust Cell Type Deconvolution (RCTD) to quantify the relative proportion of each cell-type/transcriptional state in each locale, the spot-level enrichment values for our T-cell disease factor, and spatial clusters/transcriptional niches using BayesSpace^72^ (**Fig. 6a**; **Fig S5a-j** and supplementary results).

**Figure 6.**
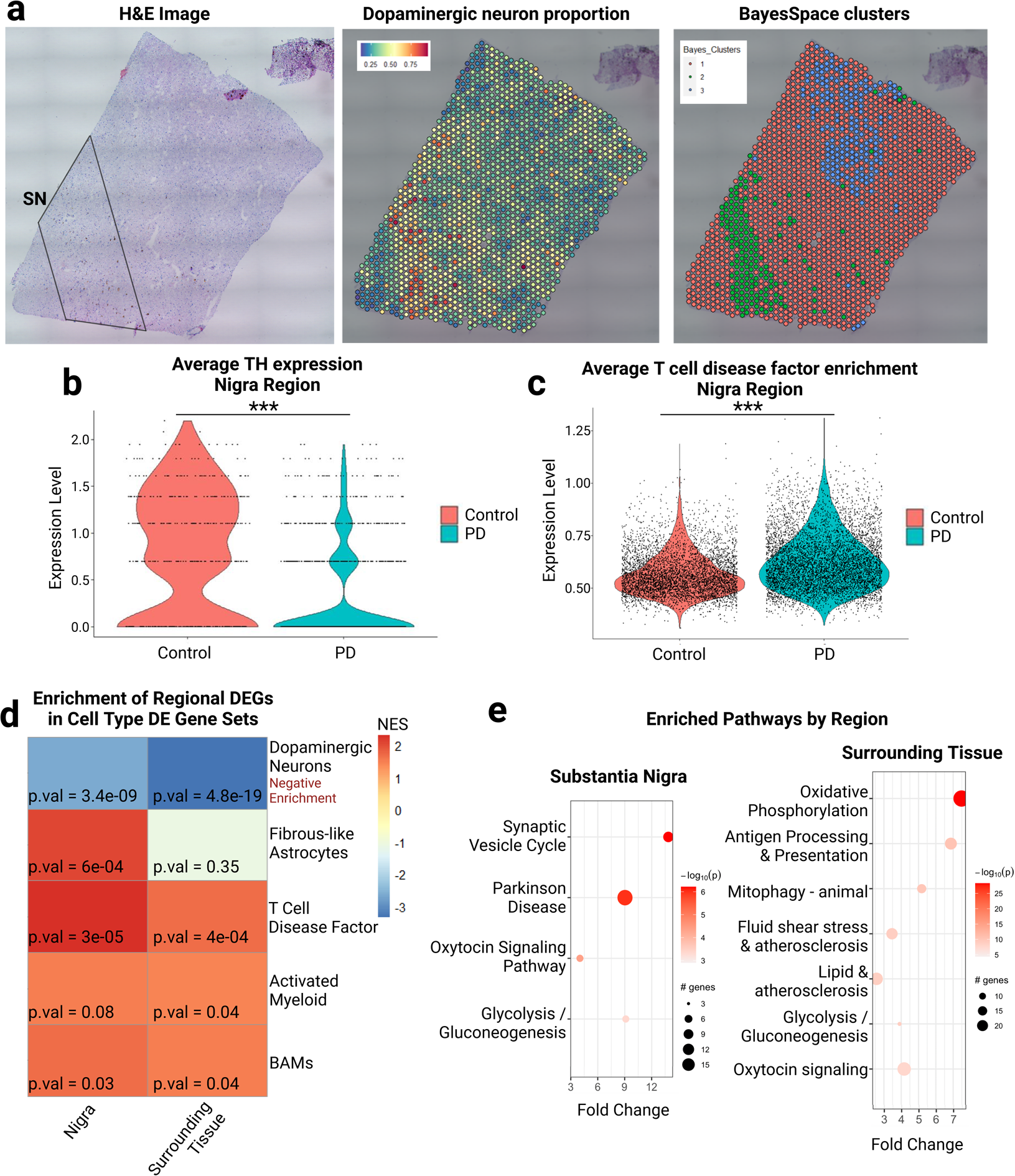
Spatial transcriptomics analyses localize cell-type specific DEGs in immune cells to local anatomic compartments in the PD substantia nigra. **a)** Example ST slide image and analysis. On the left, an H&E image of SN tissue mounted onto a 10X-Visium slide, with neurons demarcated by neuromelanin (brown). In the middle, expression values per spot of deconvolved cell type dopaminergic neurons. On the right, assigned BayesSpace clusters. **b)** Violin plot of average spot-level *TH* expression across ST tissue samples showing decreased *TH* in the PD vs control (logFC for *TH* in the SN is –1.56 with p value 6.3e-46, and in the surrounding tissue, logFC is –0.71 with p value 2.02e-94). **c)** Violin plot of average T-cell disease factor enrichment levels across PD and control ST tissue samples. The lm estimate for the surrounding tissue is –0.10, p value <2e-16. The lm estimate for the nigra in PD is 0.08, p value <2e-16. **d)** Heatmap of normalized enrichment scores from gene set enrichment analysis of the snRNAseq-derived cell-type specific DEGs. All cell types represent the increased DEGs (positive logFC), except dopaminergic neurons, which represents decreased DEGs (negative logFC). On the x axis are the two assigned regions SN and Surrounding Tissue, and on the y axis are cell types from RCTD. P values are indicated. **e)** KEGG pathway enrichment analysis in PD vs control DEGs, in SN and surrounding tissue, from ST data.

Through comparison of deconvolved dopaminergic neuron cell type proportions (**Table S1**) and *TH* expression values, complemented by neuropathological evaluation of corresponding H&E images, we annotated the BayesSpace clusters as either SN or surrounding tissue, which was predominantly white matter, including both cerebral peduncle and superior cerebellar peduncle (**Fig. S6a-k** and supplementary results). The SN region is defined by *TH* expression in addition to high proportions of dopaminergic neurons, as defined by deconvolution (see methods and **Table S1**). The surrounding white matter is defined by high proportions of oligodendrocytes (**Table S1**). As expected, *TH* expression was higher in the SN compared to surrounding white matter (regression estimate 0.24, p value <2e-16), and lower in the PD SN versus control (**Fig. 6b**). We then tested for enrichment of the T-cell disease factor in the SN, which showed significantly higher levels in PD tissue than control (**Fig. 6c**). These data independently validate our snRNAseq findings and assign the T-cell PD disease factor to the SN.

We then performed DGE analysis between PD and control ST capture areas in the two niches separately, and measured the enrichment of snRNAseq-derived cell-type specific DEGs in the ST-derived niche-specific DEGs (**Fig. 6d**; see methods). As expected, genes downregulated in PD dopaminergic neurons were decreased in both brain regions in PD. The enrichment of fibrous-like astrocyte DEGs was enriched in the DEGs of SN only, and the activated microglia/myeloid cell DEGs in the surrounding tissue DEGs only. In contrast, DEGs of monocytes/BAMs and the T-cell disease factor (factor 5 from above) were significantly enriched in both niches. This is consistent with our findings that T-cells are found in both the peduncle and in the PD SN parenchyma (**Fig. 3a-d**). We conclude that fibrous-like astrocytes adopt a disease phenotype in the SN in PD, and that monocytes/BAMs and activated microglia/myeloid cells do so in both the SN and the white matter.

Finally, we used an unbiased approach to analyze ST niche-specific DEGs and measured the enrichment of KEGG pathways in the SN and surrounding white matter (**Fig. 6e**). The results showed a marked difference in the pathways represented by each region’s DEGs. Genes involved in synaptic vesicle recycling and PD were enriched in the SN niche, which validates the results from examining TH+ neurons from the snRNAseq data. Conversely, the white matter exhibited an increase in genes involved in oxidative phosphorylation, antigen processing and presentation, and mitophagy. There were several pathways, such as oxytocin signaling and glycolysis/gluconeogenesis, that were enriched in both regions. Together, these results suggest that there are distinct, cell-type specific, spatially-defined pathologic signatures in PD.

### PD-enriched spatially defined cell-cell cohabitation and communication patterns

To determine the spatial relationships of cell types and T-cell disease factor we performed spatial cross correlation (SCC) analysis on the spot-level cell-type proportion values in our ST data (**Fig. 7a**). SCC allows us to quantify how the cell types are correlated (SCC coefficients), assign statistical significance to the coefficients using a permutation-based method, and retrieve sample-level and disease condition-level statistics. If cell types are spatially correlated, then positive SCC values will be retrieved. If they are negatively spatially correlated, negative SCC coefficients will be retrieved. Since determining SCC across thousands of ST data points is computationally intensive and slow, we developed a way to parallelize the computation, accelerating it by ∼1,200-fold (see methods and **Fig. 7b**; **Table S11**) to calculate the change in SCC between different cell-types/states observed in PD compared to control (**Fig. 7c**). The results indicate three major points.

**Figure 7.**
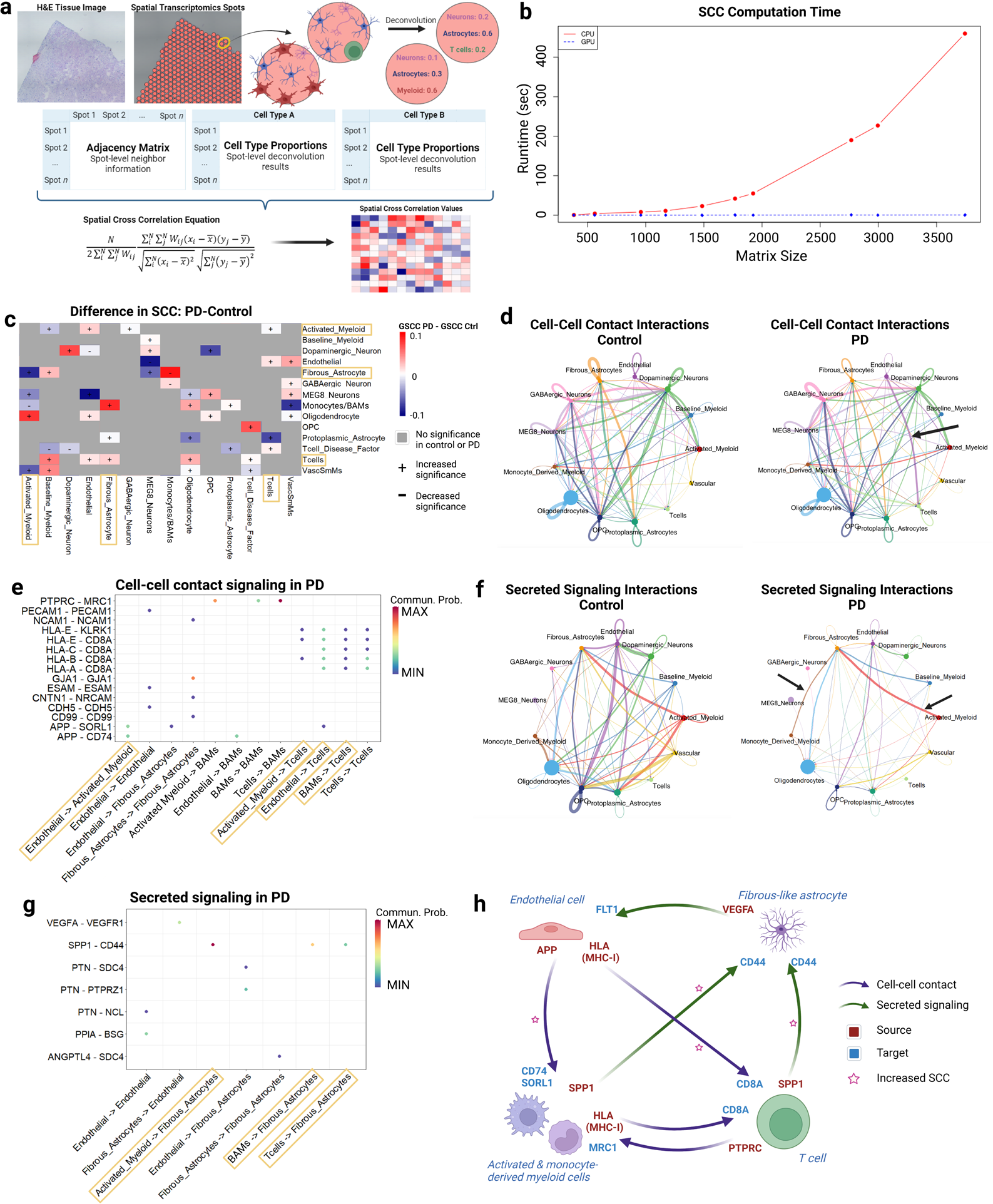
Spatial cross correlation illuminates cellular communication in PD. **a)** Schematic explaining how spatial cross correlation (SCC) values are calculated in our analyses. Spots (capture areas) are identified over tissue, and RCTD is employed to determine the cellular composition of each spot. Spot-level neighbor information is encoded in a binary adjacency matrix, which is then combined with proportion matrices for each cell type in a previously defined SCC equation. The output is a SCC value for each cell type combination. **b)** Plot of matrix size (number of elements) by amount of time (seconds) taken to complete SCC computation using our optimized algorithm conducted using the CPU (red), GPU (blue). **c)** Heatmap of change in average SCC values for each cell-type combination, PD compared to control. Increased values (red) denote an increase in SCC in PD compared to control, decreased values (blue) denote a decrease. “+” symbols represent an increase in SCC significance in PD compared to control, or a lower aggregated p value. “-” symbols represent a decrease in significance. Grayed-out boxes represent relationships that either were not significant (aggregated p value >0.05) in neither PD nor control, or that lost significance in PD compared to control. **d)** Interaction weights for SN cell types in cell-cell contact signaling, as derived from CellChat. On the left is control, on the right is PD. Note that here, “monocyte derived myeloid” refers to monocytes/BAMs. **e)** Diagram of signaling pathways implicated in cell type interactions, defined by cell-cell contact. “BAMs” refers to monocytes/BAMs, abbreviated for visual purposes. **f)** Same as c but for secreted signaling. **g)** Same as d but for secretory signaling. **h)** Schematic of the proposed potential immune-glial axis model implicated in PD, as inferred from CellChat and SCC analyses. Purple arrows represent communication cell-cell contact, and green through secreted signaling. Signaling pathways are coded with red text representing the sender, and blue text representing the recipient. Pink stars denote an increase in SCC in PD.

First, we noted increased SCC with increased significance between T-cells and oligodendrocytes, suggesting increased presence of T-cells are in the white matter. We had validated this finding by IHC for CD8 staining of postmortem PD and control tissue, not counting the cells in the subarachnoid space next to vessels (**Fig. 3a-b**), where we identified increased T-cells in the cerebral peduncle (white matter), which is rich in both cell types. Second, we found high significance in the spatial relationship between T-cells and the T-cell disease factor in PD, and lack of significance in controls. Third, we identified several cell type combinations with increased SCC, including a) activated microglia/myeloid cells and endothelial cells, b) fibrous-like astrocytes and T-cells, c) monocytes/BAMs and fibrous-like astrocytes, and d) T-cells and endothelial cells. A more detailed interpretation of SCC is provided in the supplementary results.

Taken together with our DGE results, these findings highlight novel, statistically significant patterns of increased spatial correlation and proximity between immune cells, T-cells, and other glial cells in the human postmortem PD SN.

To investigate potential patterns of altered cell-cell interactions in PD, we applied CellChat^108^, a computational method, to our snRNAseq data and identified potential interactions between pairs of cell types from inferred direct cell-cell contact and from secreted ligand-receptor signaling (**Fig. 7d-g**). We identified a potential disease-associated connection in cell-cell interaction between endothelial cells and T-cells (**Fig. 7d**), and a potential disease-associated connection in the secreted ligand-receptor communication between monocytes/BAMs and fibrous astrocytes (**Fig. 7f**).

Probabilistically, there was an increase in communication strength between endothelial cells, monocytes/BAMs, and activated microglia/myeloid cells to T-cells, characterized by MHC class I signaling to CD8A (Fig 7e). This interaction independently is consistent with the increased SCC between endothelial and activated microglia/myeloid cells on one hand and, on the other hand, T-cells. Anatomically, it is possible for this interaction to be representative of T-cells in the subarachnoid space around vessels, taking place outside of the brain.

Turning our attention to astrocytes, we noted also the increased SCCs between T-cells and fibrous-like astrocytes, and between fibrous-like astrocytes and monocytes/BAMs (**Fig. 7c**), the latter being predicted to exhibit increased secreted signaling strength (**Fig. 7f**). The end-foot processes of these astrocytes line the pial borders, allowing them to respond to soluble molecules in the cerebrospinal fluid, thus allowing them to take part in secreted signaling with cells in the subarachnoid space^109,110^. Together, our analysis suggests that, in PD, T-cells and monocytes/BAMs may communicate with fibrous-like astrocytes likely through SPP1-CD44 signaling (**Fig. 7g**).

In summary, our results outline a potential network of cell-cell interactions in PD that nominate T-cells, monocytes/BAMs and endothelial cells, along with fibrous astrocytes, as central players in facilitating neurodegeneration in the PD SN.

## Discussion

Higher levels of T-cells in PD SN than control SN and an association of PD with peripheral T-cells that recognize alpha-synuclein have been previously reported, but there has been little insight to the characteristics of these central T-cells in PD or analysis of their correspondence with peripheral T-cells. The analysis of differential gene signatures and response patterns of a plethora of immune cells, both transcriptionally and spatially, provides insights into the roles of T-cells in the PD SN.

In addition to the 831 human SN T-cells analyzed in our snRNAseq dataset, we were able to analyze over 50,000 different human T-cell clonotypes from the PD and control SN and cingulate cortex by TCR sequencing of 44 human subjects. Using independent TCR sequencing-based, computational, and *in situ* analyses, we now report that T-cells not only display a cytotoxic CD8+ tissue resident memory phenotype but are also selectively clonally expanded in the PD SN. We further propose that these T-cells have genetic and spatial profiles that indicate interactions with activated local myeloid cells, astrocytes, and epithelia within regions of the SN, and share motifs with peripheral T-cells that recognize alpha-synuclein in PD patients, despite the majority of those being CD4+ helper cell types.

In addition to the increase of clonal T-cells in the PD SN, we observed an increase in fibrous-like astrocyte genes in the substantia nigra in our spatial data, consistent with our pseudotime results, which was coupled with a an absence of upregulation of the neuroprotective protein MT3. This is similar to our previous findings in HD^38^, and highlights a region-specific response of astrocytes to neurodegeneration, which may result from, or be more likely contribute to, regional vulnerability in neurodegeneration. Additional studies are needed to determine the generalizability of this phenomenon, and whether it is a driver of resilience to neurodegeneration, a compensatory response to neurodegeneration, or both.

The spatial analyses yielded several technical and conceptual advances. We adapted our previous approach to using SCC to define patterns of cellular cohabitation in infiltrating glioma^46^ with the novel use of a new computational approach in massively-parallel GPU–accelerated fashion, that yielded computation time up to 1,200 times faster than CPU-based methods^70^ (see methods and **Fig. 7b**; **Table S11**). By leveraging the spatial data from multiple ST donors to statistically-measured changes in SCC, we constructed a spatially-informed model of cell-cell interactions in the niche of PD neurodegeneration.

Our results brought us to propose a potential immune signaling network in the PD SN that consists of interactions between astrocytes, endothelial cells, myeloid cells, and T-cells (**Fig. 7h**). We speculate that endothelial cells signal to both T-cells (HLA-CD8A) and to myeloid cells (APP-CD74), which are predicted to also signal to T-cells through cell-cell contact (HLA-CD8A). Furthermore, we found increased probability of secreted signaling from T-cells to fibrous-like astrocytes (SPP1-CD44), and from astrocytes to endothelial cells (VEGFA-FLT1), delineating an immune signaling axis through which endothelial cells mediate communication between T-cells, activated myeloid cells, monocytes/BAMs, and fibrous-like astrocytes. Finally, our analyses also highlighted increased probability of secreted signaling from activated monocytes/BAMs and T-cells to fibrous-like astrocytes (SPP1-CD44). Coupled with our spatial data, this signaling pathway is an attractive candidate to target as a means to potentially slow down neurodegeneration in PD.

Astrocyte-endothelial interaction is interesting because VEGFA-VEGFR1 signaling has been shown to increase angiogenesis, as well as microglial activation^111,112^ - which may explain the inflammatory environment in the PD SN. Moreover, activated myeloid cells participate in antigen presentation, which may drive T-cell clonal expansion^113^. In addition to antigen presentation by myeloid cells, endothelial cells can present antigens via MHC class I to cell CD8+ T-cells, which may drive activation of memory T-cells and clonal expansion^114–116^. SPP1, which is secreted by activated myeloid cells and T-cells^117,118^, interacts with CD44 and can drive downstream VEGF secretion^119,120^, potentially through the PI3K-Akt pathway^121^ (**Fig. 5e**). CD44 is mainly expressed in fibrous-like astrocytes in our dataset, and *VEGF* is differentially increased in PD SN astrocytes (**Table S4**). As a result, it is possible that SPP1-CD44 signaling to fibrous-like astrocytes may activate *VEGF*, activating *VEGFR1* in endothelial cells, perpetuating a vicious feedback cycle. This feedback cycle may lead to specific increases in local SN neuroinflammation, importantly including the replication of specific T-cell clonotypes that we have identified in PD. Future studies will attempt to validate and dissect this signaling network.

Finally, our data showed increased T-cell PTPRC signaling to monocytic MRC1. MRC1 is important for myeloid plasticity and adaptive immune responses^122^, monocytes/BAMs, and, in activated microglia/myeloid cells, can be a marker of activation^123,124^. Significantly, monocytes/BAMs (high expression levels of *CD163* and *MRC1*) have recently been shown to be necessary for an alpha-synuclein-induced neurodegeneration in a mouse model of PD^125^. The interaction between T-cells and these monocytes/BAMs via MRC1-PTPRC is consistent with previous reports^5,125^ identifying increased colocalization of both cell-types around blood vessels in postmortem PD.

Taken together, this outlines a potential immune-vascular-glial signaling axis which includes fibrous-like astrocytes, endothelial cells, myeloid cells (activated microglia and BAMs) and T-cells, and which may have the net effect of eliciting a reactive, fibrous-like state in astrocytes, activation of myeloid cells, and potentially T-cell clonal expansion in PD. Though we do not present here experimental validation for this network, it serves as a useful framework for future studies investigating the adaptive immune response in PD.

### Limitations

We note several limitations of the current study. First, our TCR sequencing data comprises only alpha-chain data: single cell TCRseq is needed to definitively identify TCR alpha-beta pairs and provide a basis for functional studies to determine precisely which antigens are recognized by specific TCR and *HLA*-antigen combinations, and whether the precise brain resident clonotypes are represented in the periphery. While this study characterizes more PD SN T-cells than previous studies, the T-cells comprise a minority of the cells and are too few to conduct single cell analysis with available technology. Furthermore, we do not have paired peripheral blood and SN samples, which will be important to define which peripheral T-cells become tissue resident after entering the CNS. Finally, as these studies are of neuropathology in subjects with advanced PD and low numbers of surviving SN dopaminergic neurons, we cannot address the issues of whether T-cells in the PD SN have increased interactions with neurons or changes in T-cell characteristics at disease stages when the highest rates of neuronal damage occur. As quantified by a board-certified neuropathologist (JEG), less than 1% (6 of 1,151) of CD8+ cells in the PD SN were observed next to neurons, discounting statistical analyses of the phenomenon.

## Author contributions

OA, KJ, JEG, DS, and PAS designed the study. OA, JLi, NM, SX, and AM performed the experiments. DC, JLee, and KJ performed processing of raw sequencing data. OA and XF collected tissue and performed dissections. KJ, OA, FP, JLee, NM, VM, PAS, JEG, and AK analyzed the data. KJ, OA, PAS, JEG, and DS wrote the manuscript and all authors read and approved the final manuscript.

## Data availability

All files, both raw and processed, used for analysis of human TCR sequencing, snRNAseq, and spatial transcriptomics datasets will be available on GEO.

All other datasets are provided in the supplementary material and/or available from the corresponding author upon request.

## Supplementary Data

### Analysis of T-cell repertoire diversity

To analyze our TCR sequencing data, we first confirmed that all repertoires were fully saturated. The saturation plots for each sample can be found in **Table S2** (see methods). Once this was established, we moved on to assessment of entropy in our repertoires.

Additional inspection of the CDR3 sequence overlap between condition and region showed that there are 2,215 global CDR3 sequences across the 44 patients in our cohort, or 14.5% of unique CDR3s in the entire repertoire (**Fig. S1a**).

To further interrogate our TCR repertoires, we quantified Shannon entropy as a measure of diversity (see methods) and partitioned the entropy to biologically known sources of diversity^39^. For each component of the T-cell receptor clonotype (**Fig. S1d**), the total entropy can be attributed to three sources: the CDR3 amino acid sequence, the VJ cassette combination, and delta between the entire clonotype and the VJ combo. For each component of the clonotype, we found that the SN exhibited significantly higher entropy compared to the cingulate, however there was no significant difference in entropy between control and PD (**Fig. S1e**). As such, we could conclude that disease condition had no bearing on the diversity of T-cell receptor repertoires, as calculated through Shannon entropy. This data is consistent with seeing more clonotypes in total in the SN versus the cingulate.

### Comparison of CDR3 Motifs to Alpha-Synuclein-Recognizing Motifs

To determine whether TCRs in the brain resemble peripheral TCRs that recognize alpha-synuclein, we used the brain repertoires as a reference in GLIPH2, and compared peripheral blood derived TCRs that recognize alpha-synuclein using a recent dataset of specific TCR sequences from CD4+ T-cells in the blood that recognize alpha-synuclein, and, as a control group, CD4+ cells that recognize pertussis toxin^7^. We identified the shared motifs and created a data frame containing a list of motifs with clonal expansion scores and contributions from alpha-synuclein-recognizing CDR3 sequences, pertussis-recognizing CDR3 sequences, and CDR3 sequences from each of our four conditions (**Table S3)**. Interestingly, there were no motifs shared by pertussis-recognizing sequences and sequences in our dataset. In contrast, 2,267 global motifs were shared by peripheral T-cell alpha-synuclein-recognizing sequences and brain T-cells. Of those, 1,204 sequences were shared with control SN, 1,087 with PD SN, 541 with control cingulate, and 849 with PD cingulate sequences (**Fig. S1h**).

Because the majority of tags with alpha-synuclein-recognizing sequences were shared with the SN, we then analyzed the differences in clonal expansion between PD and control in the SN within these shared tags. We again assigned a representative condition to each tag based on the condition contributing the highest proportion of reads, and compared the distribution of clonal expansion scores of these tags. The distributions of scores in the control group exhibited a trimodal distribution, while the PD group exhibited a bimodal distribution that was significantly shifted towards higher clonal expansion score values (Kolmogorov-Smirnov test D value 0.08, p value 0.006) (**Fig. S1i**), suggesting that while there are more T-cells present in PD SN, a higher fraction of the T-cells in the PD SN have alpha subunit motifs that share features with known TCR subunits that recognize alpha-synuclein. We note that while many TCR alpha chains are shared between CD4+ and CD8+ T-cells, identical alpha/beta combinations are rare^126^, and direct proof of specific PD SN T-cells recognizing alpha-synuclein or possessing the same TCR genes in both an individual’s brain and blood is not currently technically feasible.

### Single nucleus RNAseq identifies sublineages of neurons in the substantia nigra and cingulate cortex

As further characterization of the snRNAseq dataset, we note that from the cingulate cortex, a total of 61,870 cells passed our QC, and in the SN, 207,859 cells. The UMAP projections of these cells, grouped by donor and sex, can be seen in **Fig. S2a** for the SN and **Fig. S2e** for the cingulate. Donor and sex both evenly distributed throughout all UMAP clusters and are therefore corrected for in our batch correction approach. The number in each cell type lineage (both broad lineages, e.g. astrocytes, and sublineages, e.g. fibrous-like and protoplasmic), as well as the number of nuclei assigned to each cell type from each sample, can be found in **Table S1**.

We next analyzed the neurons from our substantia nigra dataset and found that, as expected, dopaminergic neurons clustered separately (**Fig. S2b**), and these neurons were defined by *TH* and *ALDH1A1* expression. We also found GABAergic GAD1/2+ interneurons and MEG3+MEG8+ neurons, which, unlike dopaminergic neurons, express very low *SLC6A3*, *ROBO2*, and *CALB2* (**Fig. S2c; Table S5**). When examining the DEGs across neuronal clusters, as expected, the highest number of DEGs were seen in dopaminergic neurons of the SN (**Fig. S2d**).

Next, we subclustered neurons of the cingulate cortex as we have done for the SN. Again, we found neither donor-nor sex-specific clusters (**Fig. S2e**). Sub-clusters were roughly equally represented in both conditions (**Fig. S2f**). A select subset of markers for each of the subclusters are shown in **Fig. S2g** and provided in **Table S5**. Similar to our previous findings analyzing neurons in the control and HD cingulate cortex^24^, we found layer-specific projection neurons and several inter-neuronal subtypes. Interestingly, we found that the highest number of DEGs were found in layer 2 CUX2+ glutamatergic neurons. A large number of DEGs were also noted in SEMA3E+ layer 5/6 neurons, which is expected given that Lewy bodies accumulate more in deep layers^127,128^ (**Fig. S2h**). Spatial transcriptomic studies are needed to further investigate the relationship between Lewy body accumulation and gene expression changes in PD.

### DGE analysis highlights similarities in GABAergic neurons of the cingulate cortex and substantia nigra in PD

When examining the DEGs in nigral neurons, we focused on dopaminergic neurons. Consistent with previous work^129^, we found that several of the dopaminergic neuron DEGs we involved in oxidative phosphorylation, neurodegenerative diseases, lysosome, and protein processing (**Fig. S3a**). They also showed enrichment in antigen processing and presentation^130,131^. Next, we examined the DEGs in cingulate neurons, focusing particularly on layer 2/3 CUX2+ neurons. We found that the cortical CUX2+ neurons were also enriched in pathways of neurodegeneration, as well as ErbB and Wnt signaling, ubiquitin mediated proteolysis, and endocytosis (**Fig. S3b**).

We compared the genes found to be upregulated and downregulated in PD in cortical CUX2+ neurons and dopaminergic neurons from the SN (**Tables S4** and **S5**). We found a minority of increased DEGs were shared between increased DEGs in these cell types. However, more than half of the dopaminergic neurons downregulated DEGs were shared with those of layer 2 CUX2+ cortical neurons (**Fig. S3c; Table S9)**. As such, we analyzed which pathways were enriched in these DEGs using EnrichR and its Appyter extension (**Fig. S3d**). We found that dopaminergic and CUX2+ neurons downregulated processes involved in synaptic vesicle cycle, calcium reabsorption, glycolysis/gluconeogenesis, and neurodegenerative diseases. Together, our data demonstrates that not only does PD pathology affect dopaminergic neurons in the nigra, but also layer 2 as well as deeper layer projection neurons in the cingulate cortex. That said, we do not delve into the intricacies of the neuronal pathology in this study because our focus is on the glia-immune interaction axis.

### Comparison of DEGs in Nigral Myeloid Cells

The UpSet plots in **Fig. 4g-h** indicated a total of three upregulated genes shared between nigral activated myeloid cells and monocytes/BAMs in PD: *SAT1*, *TFRC*, and *NAMPT*. The plots indicated that these genes were not increased in any other myeloid cell type of the nigra or cingulate cortex. We saw significant p values for all three genes in the activated myeloid cells and monocytes/BAMs of the nigra, however the logFC values were highest in monocytes/BAMs (**Tables S4** and **S5**). Interestingly, SAT1 has been implicated by previous studies in PD pathogenesis, exerting neuroprotective effects on brains affected by alpha-synuclein toxicity^132,133^. *TFR1*, encoded by *TFRC*, is upregulated in myeloid cells in response to inflammatory pathway signaling such as NF-kB and *HIF-1*^134,135^. Finally, *NAMPT* has also been implicated in neuroinflammation, as it is upregulated in response to inflammatory stimuli^136–138^. This data further implicates both activated myeloid cells and monocytes/BAMs in the substantia nigra in neuroinflammation and progression of PD.

### Qualification and Classification of ST Data and BayesSpace Clusters

As a quality control step, we confirmed adequate numbers of counts per spot in each tissue sample before moving on to downstream analyses. It can also be seen that areas with high numbers of counts correspond to SN tissue (**Fig. S5a-j**), consistent with the presence of neurons in these regions, as neurons express a higher number of genes compared to glial cells. Each sample was also assigned a number of BayesSpace clusters according to their qTune metrics (**Fig. S5a-j**; see methods). We asked if the clusters across different samples can be correlated, and/or anatomically annotated. Thus, we performed correlation analysis of the gene expression in each of the BayesSpace clusters (see methods), and identified three cross-sample spatial meta-clusters (**Fig. S6k**). The gene markers for the meta-clusters are provided in **Table S8**. Examining these gene markers allowed us to classify the three meta-clusters as: white matter, SN, and white matter with high expression of ribosomal genes (**Fig. S6k**). Because the white matter could not be reliably divided into specific, consistent, canonical regions, we grouped the two white matter containing meta-clusters into one category (Surrounding_Tissue) (**Fig. S6a-j**). We used these classifications for downstream analyses.

### Interpretation of Spatial Cross Correlation Data

SCC is different from traditional Pearson correlation. The diagonal of the SCC heatmap is the autocorrelation of each cell-type. It is notable that the heatmap shown in **Fig. 7c** is not symmetric along the diagonal. This is because we are showing the difference in SCC between PD and controls, aggregated across multiple samples. Also, because the relative abundance of features or cell types in this spatial dataset is spatially variable, this leads to slightly different coefficients when comparing the SCC between cell-type A and B versus SCC of the inverse relationship. In other words, the relationship of cell-type A to cell-type B is not entirely equivalent to the inverse relationship of cell-type B : cell-type A. This is because the weighting variable Wij (**Fig. 7a**), is either 1 or 0 based on the proximity of cell A to B, which is determined by the spatial abundance of each cell type relative to the other. As shown in **Fig. 7c**, the relationship between T-cells and oligodendrocytes is not equal when viewed from different axes. This can be interpreted as follows: T-cells, which are sparse cells, were colocalized with oligodendrocytes, which are abundant cells, and were rarely found where oligodendrocytes were low. Thus, the SCC of T-cell to oligodendrocytes is positive. However, The SCC of oligodendrocytes to T-cells is not equivalent. This is because the oligodendrocytes are present in many areas where T-cells are not present.

Changes in SCC relationships along the diagonal (autocorrelation) are useful to interpret, especially for sparse cells. In the SN, the TH+ neurons are evenly dispersed in the control tissue. In PD, these cells are depleted and are less abundant, thus, they become less disperse and more clustered in PD, especially given the known nigral region vulnerability of lateral vs medial tiers of SN TH+ neurons^139^. Namely, TH+ neurons are less abundant in the lateral SN and relatively less depleted in the medial SN. This neuropathologic phenomenon explains why TH+ neurons display increased autocorrelation (SCC along the diagonal) in PD, i.e., become less dispersed and more clustered, and therefore more autocorrelated. On the other hand, oligodendrocytes are present within both the SN and the white matter, the distribution of oligodendrocytes was not apparently altered in our dataset, and thus, the autocorrelation was not altered. Likewise, the spatial distribution of oligodendrocytes relative to fibrous-like astrocytes was not altered. This includes comparing the abundance of oligodendrocytes to fibrous-like astrocytes and vice versa. This is because both cell-types are more abundant in the white matter, which represents a large fraction of our ST dataset.

## Supplementary Figure Legends

**Figure S1: TCR entropy and a-syn GLIPH2 results**

**a)** Venn diagram of all CDR3 sequences in the TCR repertoires, across all patients, grouped by region and condition. On the bottom left (purple oval) is the cingulate cortex control, top left (yellow oval) SN control, top right (green oval) SN PD, and bottom right (pink oval) cingulate cortex PD. **b)** Ridge plot showing the abundance distributions corresponding to each region and condition in the global CDR3s (center section in Venn diagram). The log-normalized reads are on the x axis, and region and condition are on the y axis. The left side represents sequences with higher numbers of reads, the right side lower numbers of reads. **c)** Bar plot depicting the average number of CDR3 sequences seen in more than one patient in control (orange) and PD (blue) samples of the substantia nigra. **d)** Entropy values for clonotype components - For each component, we found that the substantia nigra exhibited significantly higher entropy (linear model p values 0.0237, 0.024, 0.0131, and 0.0464, respectively), however there was no significant difference in entropy between control and PD (linear model p values 0.4487, 0.537, 0.3775, and 0.4010, respectively). **e)** Bar plot of p values from comparison of entropy for each clonotype component, measured by linear regression between condition (orange bars) and region (blue bars). The y axis represents the calculated p value, while the x axis shows the clonotype component being compared. **f)** Box plots depicting clonal expansion scores for each of our four conditions, calculated from the patient level. The y axis represents D values derived from Kolmogorov-Smirnov testing. Values between control and PD in the cingulate cortex were not significantly different (p value 0.83). SN control D scores were significantly lower than SN PD, with p value 0.048. **g)** Plot of significance distributions generated 100-iterations from patient-level clonal expansion scores (see methods). The x axis shows the generated p values, and the y axis shows the density. Values from the SN are represented in blue, cortex in orange. **h)** Pie chart of the number of GLIPH2 tags/motifs shared between the alpha-synuclein reactive peripheral T-cells and each of the four specified conditions from our dataset. **i)** Ridge plot of clonal expansion scores of GLIPH2 tags/motifs shared between the alpha-synuclein reactive peripheral T-cells and control and PD SN. **j)** Gene set enrichment of a general memory T-cell geneset in T-cell gene expression in PD. Normalized enrichment score is 2.01 with p value 7e-05.

**Figure S2: Single Nucleus Analysis of Neurons in the Substantia Nigra.**

**a)** UMAP plots of all substantia nigra nuclei, grouped by donor on the left, and by sex on the right. **b)** UMAP plots of substantia nigra neuronal nuclei, grouped by sublineage on the left, and by condition on the right. **c)** Dot plot of select marker genes for each neuronal sublineage in the SN. Color represents the average normalized gene expression value, and size represents the percentage of cells expressing the gene. **d)** Number of differentially expressed genes in each neuronal sublineage of the SN. Green bars represent the number of genes with negative logFC values, or downregulated in PD, and blue bars represent those with positive logFC values, or upregulated in PD. **e)** Same as a but for the cingulate cortex. **f)** Same as b but for neurons in the cingulate cortex. **g)** Same as c but for neurons in the cingulate cortex. **h)** Same as d but for neurons in the cingulate cortex.

**Figure S3: Comparison of Gene Expression in Neurons and Myeloid Cells**

**a-b)** Pathway enrichment of DEGs from dopaminergic neurons in the SN (**a**) and layer 2 CUX2+ neurons in the cingulate cortex (**b**). The fold enrichment is represented on the y axis, and each pathway on the y axis. Dot color represents –log10 p values, and dot size represents the number of genes in the pathway. **c)** Venn diagrams showing the overlap of DEGs, both increased (left) and decreased (right) between dopaminergic neurons in the SN and layer 2 CUX2+ neurons in the cingulate cortex. **d)** Bar plot showing the KEGG pathways represented by the shared decreased DEGs between the SN dopaminergic neurons and the cortical layer 2 CUX2+ neurons.

**Figure S4: Single Nucleus Analysis of Astrocytes in the Cingulate Cortex.**

**a)** UMAP plots of all cingulate cortex astrocyte nuclei, grouped by sublineage (protoplasmic in orange, fibrous-like in blue), condition (control in orange, PD in blue), and donor. **b)** Number of DEGs in each astrocytic sublineage in the cingulate cortex. Green bars represent the number of genes with negative logFC values, or downregulated in PD, and blue bars represent those with positive logFC values, or upregulated in PD. **c)** KEGG pathway enrichment of DEGs in fibrous-like and protoplasmic astrocytes in the cingulate cortex. **d)** Dot plot of select markers genes for each astrocyte lineage in the cingulate cortex. Color represents the average normalized gene expression value and size represents the percentage of cells expressing the gene. **e)** Pseudotime plot of astrocytic nuclei in the cingulate cortex, with astrocytes expressing protoplasmic genes on the left side in red, and astrocytes expressing fibrous-like genes on the right in blue. Nuclei are projected on PHATE axes. Color bar indicates pseudotime value range. **f)** Ridge plot depicting the proportion of protoplasmic and fibrous-like in low to high pseudotime values. Protoplasmic astrocytes are depicted in orange, fibrous-like in blue. **g)** Ridge plot depicting the enrichment of PD and control astrocytes in low to high pseudotime values. PD astrocytes are depicted in blue, control in orange.

**Figure S5: Spatial Transcriptomics and BayesSpace Spot-Level Data.**

**a-f)** From left to right: Original H&E tissue image, number of counts per spot (normalized by SCT), and regional tissue classifications for each sample. **g-j)** same as a-f, but tissue image is stained with antibodies NeuN, GFAP, and DAPI.

**Figure S6: BayesSpace Clusters and Their Correlations.**

**a-j)** BayesSpace cluster assignments for each spot in each sample. **k)** Heatmap of correlated clusters for each sample as defined by correlation of gene expression. The large top left cluster is classified as white matter, the middle cluster as substantia nigra (SN), and the lower right clusters as white matter with increased ribosomal gene expression. The BayesSpace clusters for each sample are indicated on the x and y axes. Dark red colors denote a high correlation value, and dark blue colors denote a lower correlation value between variable feature enrichment.

**Figure S7: MT3 and GFAP staining in astrocytes**

**a-b)** Cells in the cingulate stained for DAPI (blue) to detect nuclei of all cells and GFAP (green) to detect astrocytes. Scale bar = 20 μm. **c-d)** Cells in the SN stained with DAPI (blue) to detect nuclei of all cells and GFAP (green) to detect astrocytes. Scale bar = 20 μm. The next row shows MT3 (red) alone. The last figure is the merged of all three channels. **e)** Example of “GFAP high” cells shown by arrows. **f)** Example of “GFAP low” cells shown by arrows. **g)** Quantification of the proportion of MT3 positive astrocytes in their respective regions. Unpaired two-tailed T-test N = 6 for both conditions. P value = 0.0017 for the proportion of MT3 positive GFAP-high astrocytes in the cingulate, p value = 0.9290 in the SN. The proportion of astrocytes labeled “GFAP High” in the cingulate has p value = 0.003, SN has p value = 0.0019. Data is shown as mean +/−SEM.

## Supplementary Tables

**Table S1:** T-cell Receptor Sequencing, Single Nucleus, and Spatial Transcriptomics Sample Metadata.

**Table 2:** Saturation Plots from TCR Sequencing Repertoires.

**Table 3:** Expanded CDR3 Sequences and GLIPH2 Results.

**Table 4:** Substantia Nigra Lineage Cluster Markers, DEGs, and PathfindR Results.

**Table 5:** Cingulate Cortex Lineage Cluster Markers, DEGs, and PathfindR Results.

**Table 6:** CD8+ Tissue Resident Memory Gene Set and General Memory Geneset.

**Table 7:** scHPF Cell Scores and Gene Scores.

**Table 8:** BayesSpace Cluster Correlation Data.

**Table 9:** Comparison of DEGs in Cingulate Cortex and Substantia Nigra Neurons and Myeloid Cells.

**Table 10:** Antibody Descriptions.

**Table 11:** Comparison of GPU versus CPU SCC Calculations.

## Acknowledgements

This work was supported by the Aligning Science Across Parkinsons’s (ASAP) initiative and the Michael J. Fox Foundation (MJFF).

