## Supplementary figures and images for "The spatial landscape of glial pathology and T-cell response in Parkinson’s disease substantia nigra"

### Supplementary Figures

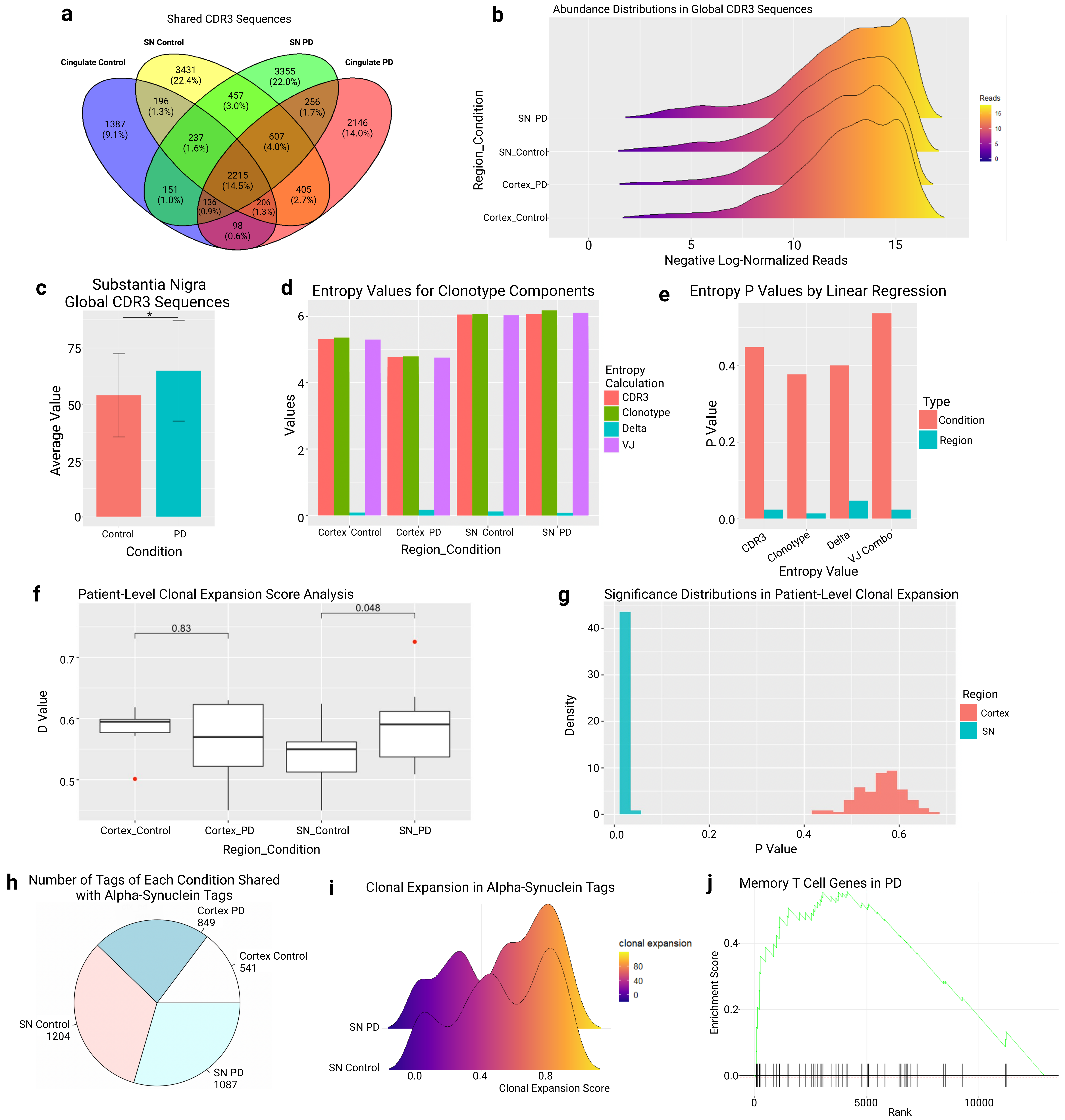

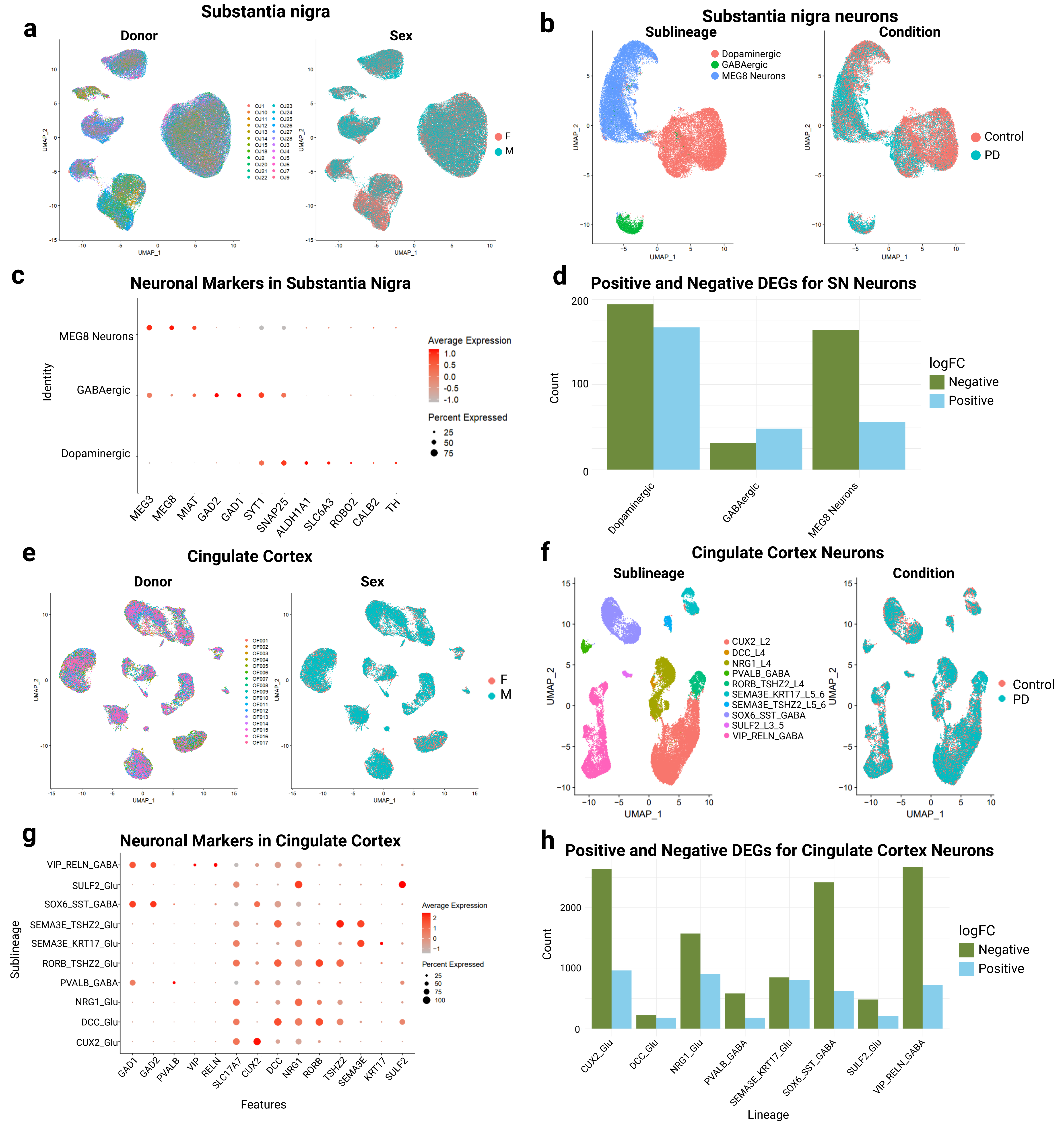

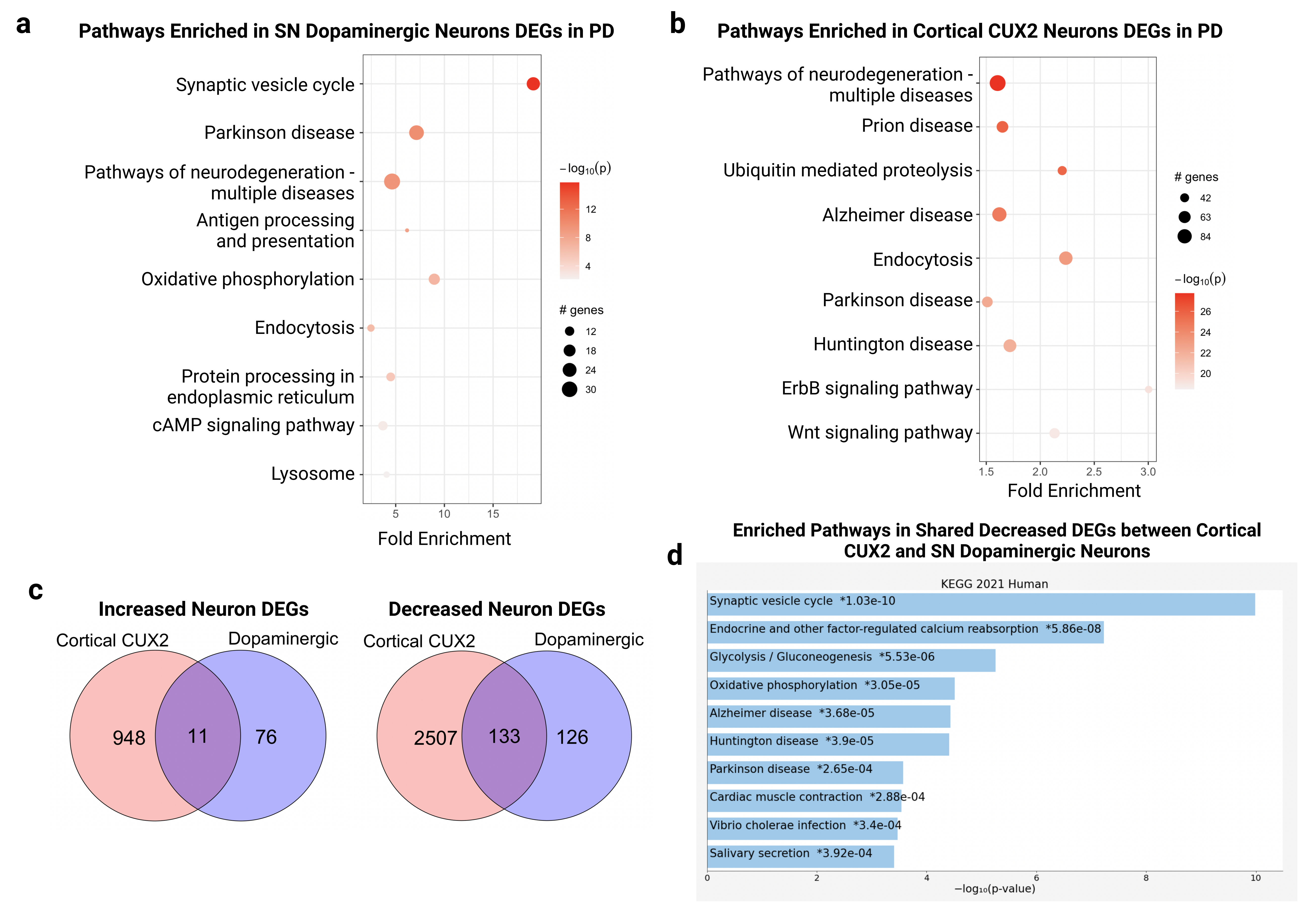

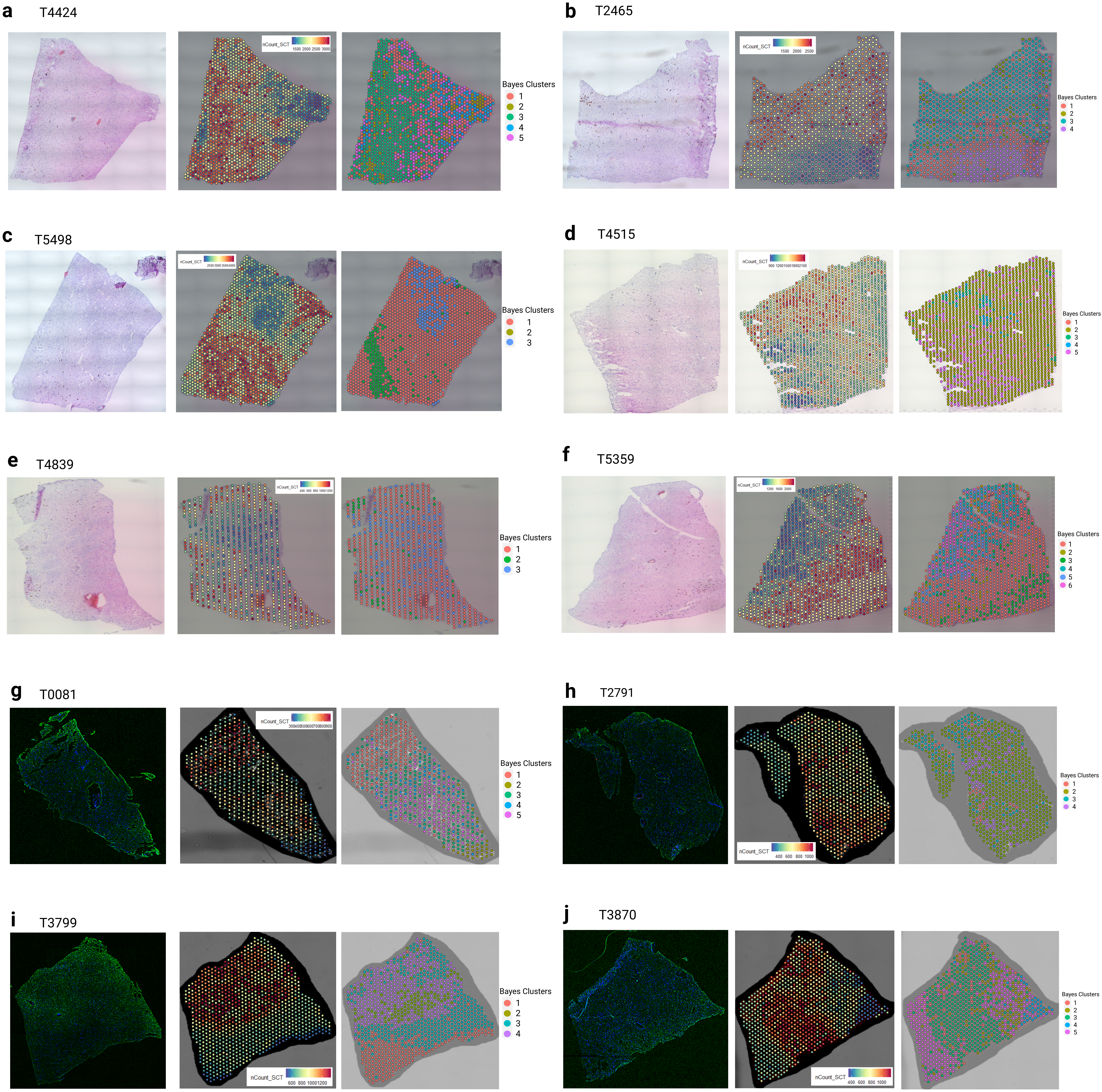

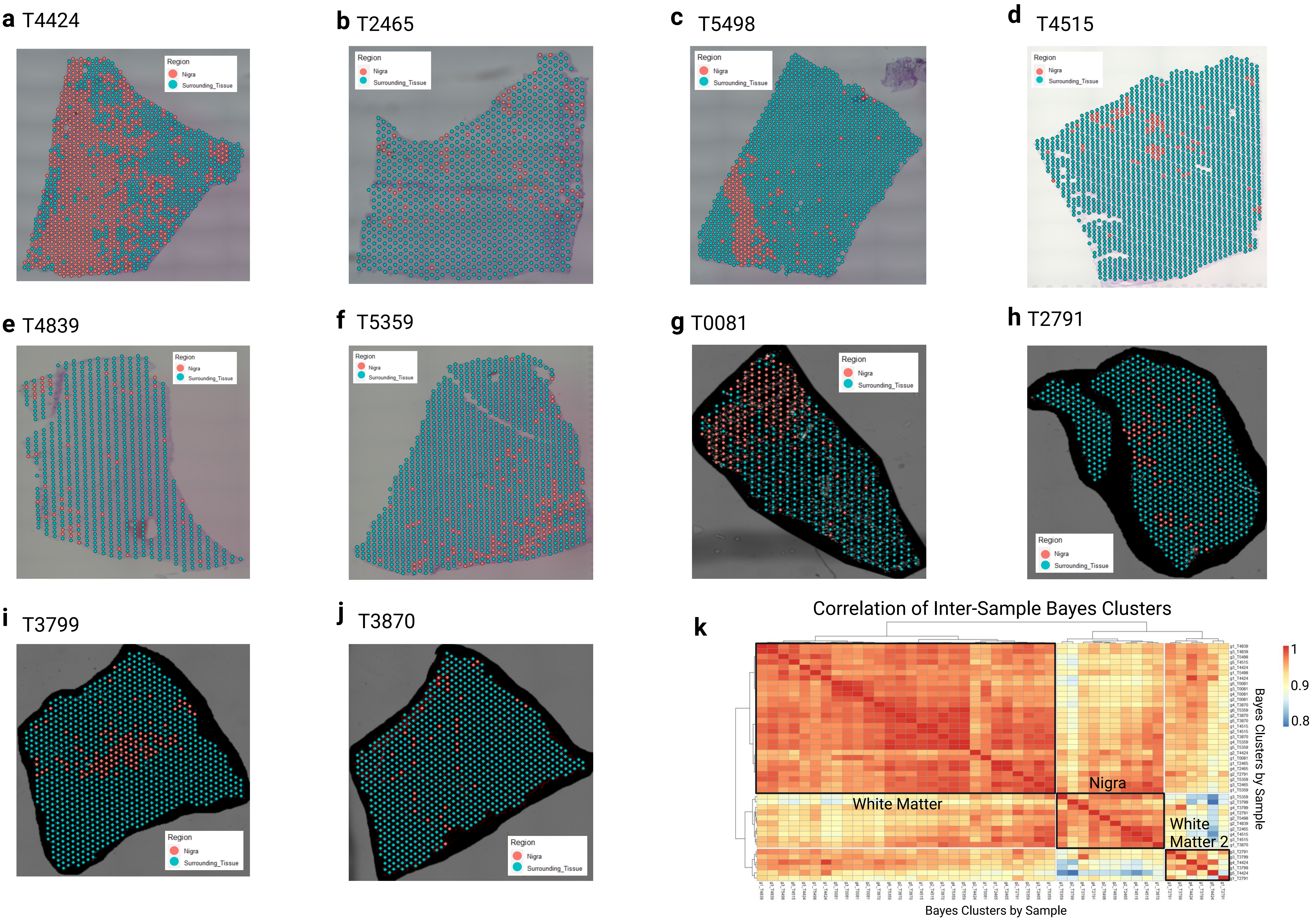

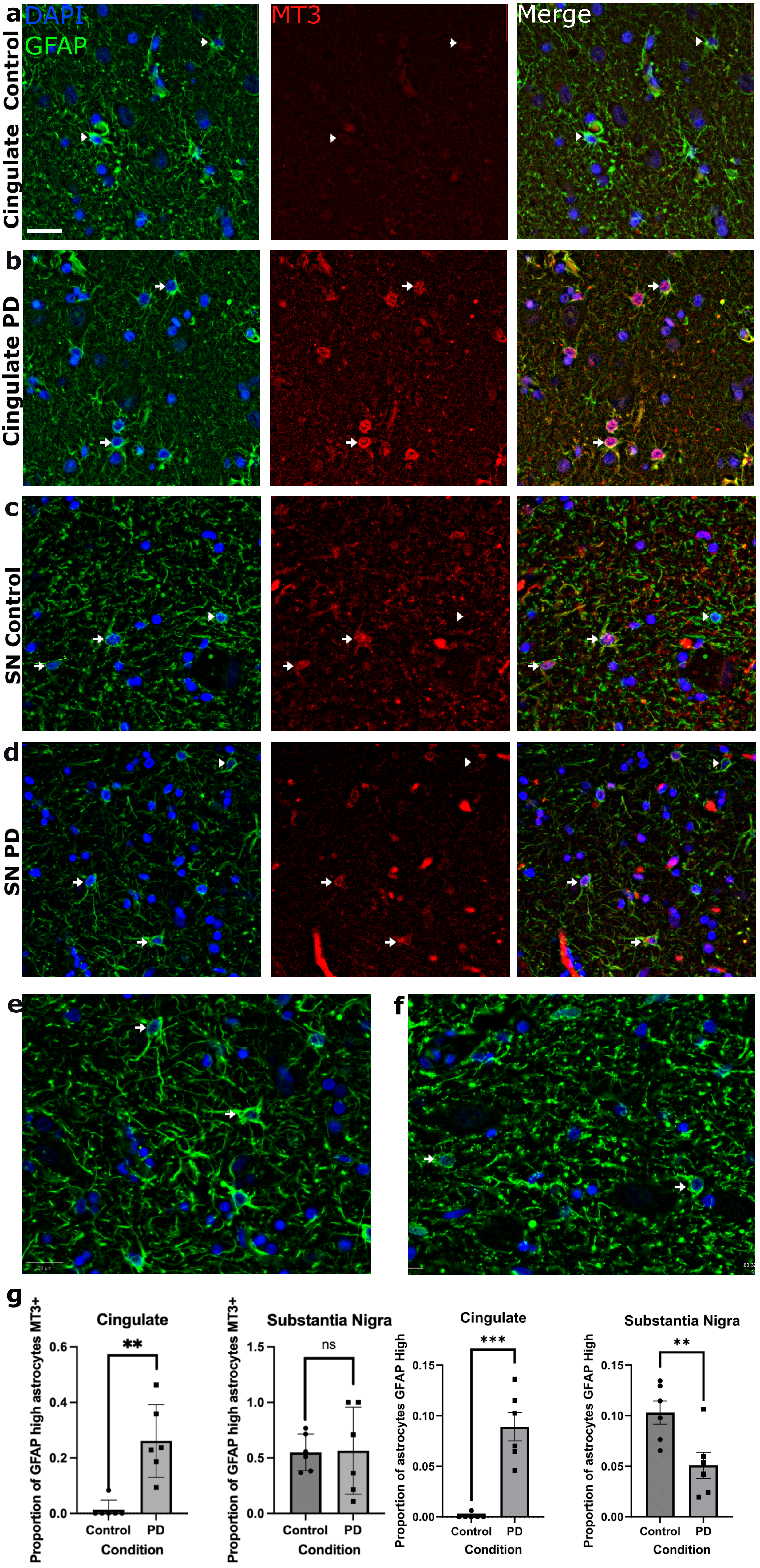
